# Human brain cell types shape host-rabies virus transcriptional interactions revealing a preexisting pro-viral astrocyte subpopulation

**DOI:** 10.1101/2024.12.18.629263

**Authors:** Lena Feige, Karl Young, Juan Pablo Cerapio, Tatsuya Kozaki, Lauriane Kergoat, Valentina Libri, Florent Ginhoux, Milena Hasan, Lamya Ben Ameur, Gregory Chin, Zach Goode, Hervé Bourhy, Arpiar Saunders

## Abstract

How virus-host cell interactions and innate immune antagonism shape neurotropic infection dynamics across diverse brain cell types are largely unknown. To “unmask” and study how innate immune inhibition affects cell type-specific transcriptional regulation of the human and viral genome, we performed single-cell RNA sequencing of human brain cell co-cultures, comparing an isolate of rabies virus (RABV) to its mutant incapable of antagonizing interferon- and NF-κB-dependent responses. RABV gene expression was shaped by host cell type. RABV induced small-scale, cell-type conserved transcriptional programs that likely support infection by 1) hijacking negative transcriptional feedback of pro-viral factors while 2) reducing anti-viral RNAs. Unexpectedly, disinhibited innate immune signaling increased RABV transcription, most strikingly in an infection-independent “pro-viral” astrocyte subpopulation. Further analysis suggested that “pro-viral”-like astrocytes are a rare subtype in the human brain and are primed to protect the brain during viral infection in concert with interferon-sensitive microglia recalcitrant to infection.

## Introduction

Viruses possess diverse mechanisms that shape the cellular and molecular properties of host cells through direct viral-host protein interactions. Due to historic technical limitations, the molecular consequences of these interactions have been difficult to characterize, especially across diverse primary cell types that constitute complex tissue such as the human central nervous system (CNS). The neurotropic Rabies virus (RABV) – a prototypic member of the negative-stranded RNA *Lyssavirus* genus – has evolved specialized mechanisms of immune evasion in the CNS, contributing to fatal encephalomyelitis and approximately 60,000 deaths annually^1^. Immune evasion is a critical feature of RABV pathogenesis, as the virus must travel undetected for long distances from peripheral muscle into the CNS, leveraging neurons and neuronal synapses for retrograde (dendrite-to-axon) transport^2^. Remarkably, neuronal viability and function are preserved during infection of non-cell culture adapted RABV strains as brains often show minimal macroscopic and histopathological alterations^3,4^. While neurons possess mechanisms to detect and respond to infection, neuronal immune responses are comparatively restricted to those of neighboring glia^5,6^, functionally-distinct brain cell types that form intimate physical interactions with neuronal processes and perform myriad functions. Recent *in vivo* evidence suggests RABV can infect astrocytes and Schwann cells, challenging the notion of exclusive neuron-specific tropism^7–9^. Astrocytes in particular are crucial for IFN-β signaling during RABV infection^10^ and can enter “reactive” molecular states in response to diverse insults, including viral infection^11^. Microglia too are involved in RABV pathogenesis^12^, but it remains unclear whether viral antigens observed within microglia are due to active infection or because of phagocytosis of infected neuronal material. Despite this unresolved debate, numerous reports indicate microglia transcribe genes involved in innate immunity and signaling (such as *ISG15*, *OASL1*, *IFIT2*, *CCL5*, *CXCL10* and *IL-6*) in response to RABV infection^9,13,14^. Whether these transcriptional responses are driven by direct microglia infection versus cytokine signaling from other cell classes remains unclear. To counteract cellular innate immune pathways, RABV proteins evolved mechanisms to sabotage the *nuclear factor ‘kappa-light-chain-enhancer’ of activated B-cells* (NF-κB)^15,16^ and the Janus kinases (JAK) - Signal Transducers and Activators of Transcription (STAT) pathways^17–20^, and thus potently inhibit host cell transcriptional responses downstream of interferon (IFN) type I signaling^21^. Although significant mechanistic insights have been made into immune inhibitory function of RABV proteins using cell lines, the consequences of these physical interactions on properties of infection in neurons and glia are largely unknown. Importantly, many viruses, including RABV, appear to have co-opted host cell proteins (including innate immune detectors^22^) for their life cycle^23^, suggesting that infection outcomes may depend on a complex relationship between anti- and pro-viral factors^24^, the nature of which remains to be fully elucidated.

In this study, we developed an experimental system using Tha isolate RABV, co-cultured stem cell-derived human brain cell types and single-cell RNA-sequencing (scRNA-seq) to discover how active inhibition of IFN and NF-κB signaling shapes transcriptional responses across the human and viral genomes at single-cell resolution. RABV genes are transcribed by the RNA-dependent RNA polymerase L and the phosphoprotein P complex at levels that reflect the order of the negative stranded RNA genome (3′ N-P-M-G-L 5’)^25–27^(**Figure S1A,B**). The influence of cell type and state on RABV mRNA levels is unknown.

Of the five RABV proteins, the phosphoprotein (P) and matrix protein (M) potently inhibit key pinch-points in cellular innate immune signaling. Specifically, P protein selectively binds and inhibits phosphorylated targets downstream of IFN signaling, including STAT1/2/3^17–19,28–30^ transcription factors in the cytoplasm or nucleus, as well as TBK1 and IKKε kinase^31–33^. M protein binds and inhibits RelAp43^15,16^ - an N-terminally truncated isoform of the RelA NF-κB transcription factor - disturbing NF-κB transcriptional complexes (**Figure S1C**). Foundational work has shown that 1) RABV adaptation to cell culture genetically erodes the potency of innate immune inhibition^21^ and alters viral tropism *in vivo*^7^ and 2) has identified mutations in outward facing residues in P (W265G, M287V; “2P”)^34^ and M proteins (R77K, D100A, A104S, M110L; “4M”)^15^ that prevent pSTAT1/2 and RelAp43 binding, respectively, yet leave canonical viral functions intact^20^ (**Figure S1D-F**). In response to exogenous IFNα stimulation, Tha RABV completely blocks activation of transcriptional activity of the interferon-stimulated gene (ISG) response element. However, when Tha2P4M mutations are present, transcriptional activation of the ISG response element is equivalent to uninfected cells (i.e. fully disinhibited)^20^. Thus, we sought to “unmask” the otherwise potent innate immune inhibition of Tha virus using its 2P4M knock-in mutation counterpart and study the consequences of intact and unmasked innate immunity on viral and host transcriptional signatures across human brain cell types.

Our analysis reveals several important discoveries. First, we show that Tha RABV gene transcription is coupled to transcription of the host cell genome and shaped by the host cell type. Second, we demonstrate that Tha virus induces a conserved transcriptional response in both neurons and astrocytes, increasing RNAs encoding proteins that are likely “pro-viral” and thus promoting infection, while decreasing RNAs encoding interferon-related transcription factors that control anti-viral responses. Third, we show, counterintuitively, that 2P4M-based disinhibition of innate immune signaling is associated with an increase in RABV transcription across cell types. Increased viral transcription is most strikingly observed in a subpopulation of “pro-viral” astrocytes, likely enabled by large-scale changes in the astrocytic molecular milieu. These changes include increased transcription of NF-κB effector and interferon-stimulated genes (ISGs) and reduced expression of genes supporting protein translation and mitochondrial function. Additional analysis suggests that “pro-viral” astrocytes are likely a rare subpopulation present in nominally uninfected human brain. Finally, we demonstrate that microglia are almost completely recalcitrant to infection but nonetheless respond to infection of other cell types through an ISG expression response that is potentiated by both higher viral challenge doses and 2P4M mutations. Through the lens of gene expression, our data collectively suggest that innate immune inhibition by isolate RABV maintains infection stealth by restricting viral gene expression, inhibiting the emergence of “pro-viral” astrocyte state and limiting ISG transcription in microglia.

## Results

### Discerning infection states and host cell types from single-cell RNA sequencing data

To study how isolate Tha RABV alters the molecular properties of human brain cell types at single-cell resolution - and to “unmask” and then characterize how antagonism of IFN and NF-κB by Tha RABV P- and M-proteins sculpt these changes through regulation of both the human and Tha RABV genome - we performed scRNA-seq analysis (10x Genomics Chromium 3’ v3.1) of co-cultured human neurons, astrocytes and microglia 48 hours after bath application of Tha (multiplicity of infection (MOI) 0.5 or MOI 5) or Tha2P4M (MOI 0.5) or from uninfected control wells (“No Infection”)(**Figure 1A**). Absolute MOIs were calculated using interferon-response deficient BSR cells yet, when introduced into co-cultures, represent a relative exposure of ten-folder lower or higher number of RABV particles (**Methods**). Brain cell co-cultures were seeded with approximately equal proportions of hNSC-derived neurons and astrocytes (∼47% each) as well as a small population microglia derived from human pluripotent stem cells(∼6%)(**Methods**). The presence of these differentiated cell classes and Tha RABV infection was confirmed using a Tha virus expressing eGFP followed by immunolabeling of cell-type-specific marker proteins (**Figure 1B**). After quality control, our analysis focused on n=23,457 single-cell RNA profiles derived from acutely dissociated cells with >2,000 RNA transcripts (as defined by unique molecular identifiers; UMIs) collected across multiple biological replicates (No Infection control, n=3; Tha(0.5), n=3; Tha(5), n=2; and Tha2P4M (0.5), n=2)(**Figure S2A,B; Methods**).

**Figure 1.**
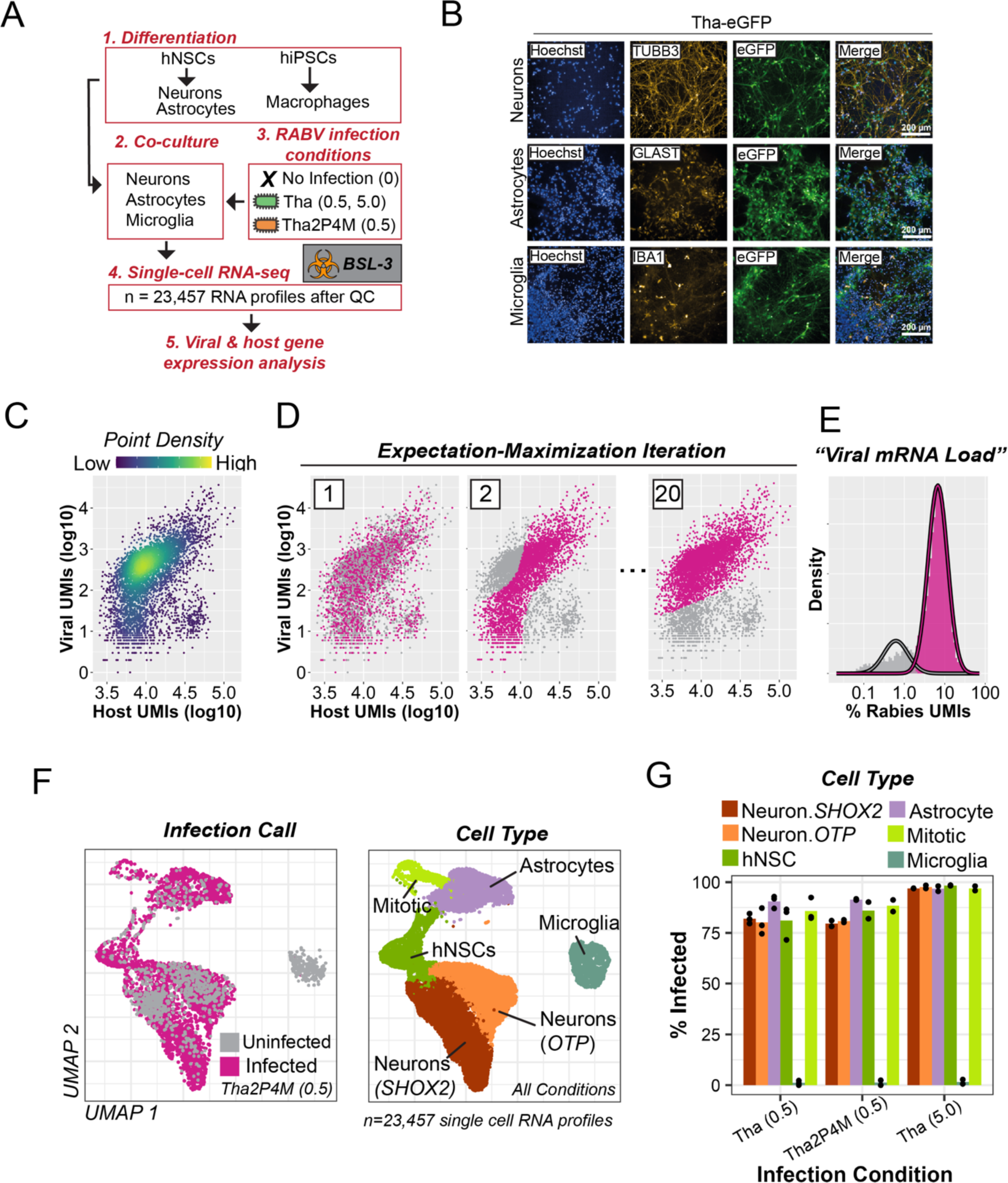
Unmasking the transcriptional effects of innate immune signaling inhibition across human brain cell types using field-isolate Tha wild-type and 2P4M RABV and single-cell RNA-seq analysis. **(A)** Overview of the study system and scRNA-seq data generation. Human induced neural stem cells (hiNSCs) or pluripotent stem cells (hiPSCs) were transformed into neurons/astrocytes or macrophages and co-cultured, further inducing macrophage transformation into microglia. Under Biohazard Safety Level 3 conditions, scRNA-seq analysis was performed on acutely dissociated cells 48 hours after no viral infection (n=3 replicates) or bath application of Tha virus at MOI 5 (Tha(5); n= 2 replicates) or MOI 0.5 Tha(0.5); n= 3 replicates) and Tha2P4M virus at MOI 0.5 (Tha2P4M(0.5); n=2 replicates). After quality control, the final dataset consisted of 23,457 RNA profiles and included 23,528 human and 5 RABV genes. **(B)** Immunolabeling of cell type marker proteins on co-cultured cells after infection of Tha RABV expressing eGFP (Tha-eGFP), confirming the presence of infected human-induced neurons (TUBB3), astrocytes (GLAST) and microglia (IBA1). (**C-E**) Transcriptional identification of infected and uninfected cell types. **(C)** Density plot comparing the number of viral UMIs versus total (viral + human) UMIs for each scRNA profile in the analysis. **(D)** Systematic identification of infected and uninfected scRNA profiles using an Expectation-Maximization modeling approach to distinguish two cell populations based on viral UMIs (regressed by total UMIs)(**Methods**). Example results of infected (magenta) and uninfected (gray) scRNA profiles at iteration 1 (random assignment), iteration 2 and 20 (after convergence) are shown. **(E)** Density plots of “viral mRNA load” (the percentage of total UMIs that are viral) for scRNA profiles called as infected (magenta) or uninfected (gray). (Infected, mean = 3.9±0.03 sem; range: 0.38 – 64.8; Bystander, mean = 0.17± 0.002 sem; range: 0-0.64). **(F)** UMAP plot showing two dimensional embeddings of scRNA profiles following Seurat-based clustering analysis using exclusively human genes (**Methods**). scRNA profiles colored by infection status (*left*) or cell type-based cluster assignments (*right*). **(G)** Barplots of the percent infected cells stratified by experimental condition and cell type.

We next sought to accurately distinguish infected cells and uninfected “bystander” cells and assign host cell type identities to each RNA profile. Because infected cells may contain variable levels of viral mRNA, and bystander cells may accrue viral transcripts by chance during cell dissociation and scRNA-seq library preparation, we developed a data-driven computational approach using expectation-maximization to iteratively refine viral RNA distributions for infected and bystander populations, accounting for variability in viral RNAs driven by biological and technical factors (**Figure 1C,D**; **Methods**). The resulting “viral mRNA load” distributions – defined as the percentage of total cellular mRNA derived from the Tha genome – clearly delineated populations of infected and bystander cells (**Figure 1E**). Simulated viral expression data based on these inferred distributions closely matched that of real data, confirming the accuracy of our approach (**Figure S2C**). Throughout this study, we use both qualitative infection categories as well as a quantitative metric of infection progression (“viral mRNA load”) to guide our analysis of host and virus gene expression relationships at single-cell resolution.

To assign single-cell RNA profiles to transcriptionally-defined brain cell types, we clustered cell profiles using only genes from the human genome (**Methods**). The resulting clusters corresponded to 5 major brain cell populations: neural stem cells (10.7%; n=2,526), mitotic cells 3.3%; n=781), astrocytes (16.9%; n=3,975), microglia (5.2%; n=1,218) and two groups of neurons (*OTP*+ and *SHOX2*+, (31.2%; n=7,312 and 32.6%; n=7,645, respectively) (**Figure 1F** and **Figure S2D**). Each clustered cell population was robustly delineated by hundreds of differentially expressed genes, including well-established “markers” that confirmed cell type identities (**Figure S2E**). Throughout our analysis, we used both OTP+ and SHOX2+ populations when analyzing neurons. RNA expression relationships were consistent with a dynamic developmental continuum visible in the two-dimensional UMAP embedding, with a population of hNSCs differentiating into both neurons and astrocytes, as well as a population of markedly distinct microglia, consistent with unique developmental and experimental origin.

We next evaluated the type and proportion of infected cells across cell types and Tha RABV infection conditions (**Figure 1G**). While microglia exhibited strikingly low infection percentages across all conditions (∼1%), all other cell types exhibited infection percentages between 79.7-91.1% at MOI 0.5 and 96.8-98.4% at MOI 5. We observed no difference in cell-type rate of infection between Tha(0.5) and Tha2P4M(0.5), suggesting that mutations in the RABV P- and M-proteins had little effect on infection probability. Collectively, our scRNA-seq data and analysis framework confirm the tenets of our human cell type co-culture experimental system and enable properties of Tha infection and the consequences of innate immune inhibition to be analyzed at single-cell resolution in the context of host cell type.

### Tha2P4M mutations increase viral transcription

We next sought to understand how RABV transcription is shaped by human host cell type and the 2P4M mutations. Because lyssavirus infection involves both negative-stranded RNA genomes and positive-stranded anti-genomes and transcripts, we first evaluated the genomic locations of our Tha RABV scRNA-seq reads to determine which RNA species were counted in our dataset. Consistent with exclusive mRNA capture and 3’ RNA sequencing, we observed sense-read pileups immediately adjacent to viral polyadenylation signals, with negligible sense or anti-sense reads mapped to other genomic locations. As expected, reads from bulk sequencing of whole Tha mRNAs^35^ fully covered the five RABV gene bodies (**Figure S3A,B**). Next, we evaluated mRNA-based viral load distributions across cell types and experimental conditions (**Figure 2A** and **Figure S3C**). We observed that mRNA viral load scores did vary by both cell type and experimental condition. Strikingly, Tha2P4M(0.5) infections exhibited a robust, consistent doubling of viral mRNA load across cell types (range of means = 4.8 – 7.7%) as compared to Tha(0.5) and Tha(5) infections consistently (range of means: Tha(0.5), 3.3 – 3.8%; Tha(5), 3.2 – 3.8%. Since the 2P4M mutations do not detectably alter the function of RABV transcription or replication^20^, these results suggest cellular changes downstream of the 2P4M mutations (likely through disinhibited IFN and NF-κB signaling) increase rather than restrict viral transcription across cell types. Moreover, a subset of Tha2P4M infected astrocytes exhibited strikingly high viral mRNA load scores (>10%) not observed in other Tha2P4M infected cell types (**Figure 2A**), suggesting 2P4M-disinhibited RABV transcription is especially pronounced in a subpopulation of human astrocytes. Thus one role of wild-type Tha P and M proteins is to indirectly restrict viral mRNA expression as compared to 2P4M mutated proteins.

**Figure 2.**
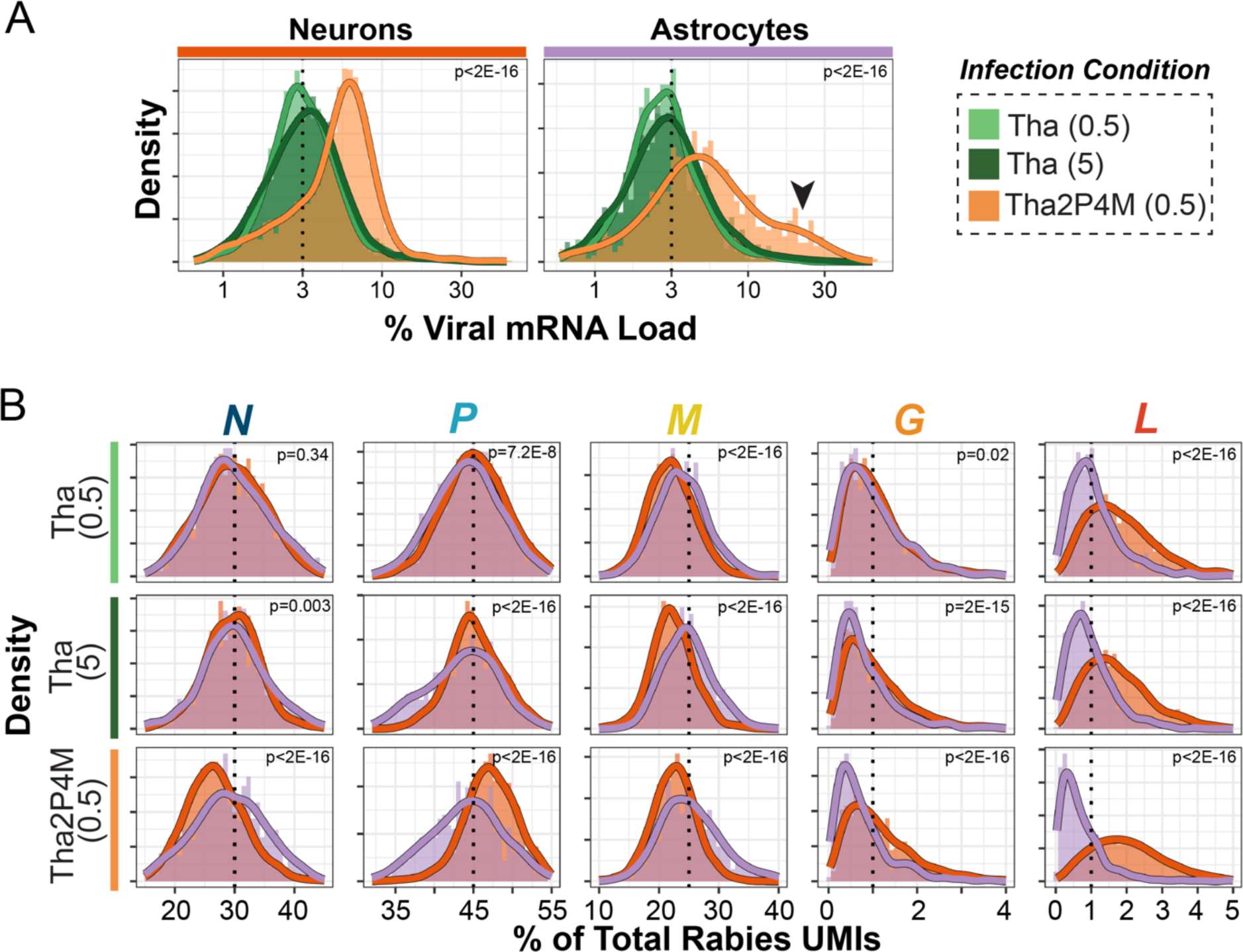
Tha2P4M mutations double viral mRNA load without major alterations to cell-type-specific viral gene expression. **(A)** Combined density and histogram plots of viral mRNA load stratified by neurons and astrocytes and color-coded by experimental condition (n cells: Neurons, Tha(0.05) = 2,711; Tha(5) = 1,877; Tha2P4M(0.05) = 2,080; Astrocytes, Tha(0.05) = 1,036; Tha(5) = 798; Tha2P4M(0.05) = 806). Arrowhead indicates high viral mRNA load astrocytes. Both cell type and experimental condition explain differences in viral load distributions (p<2E-16, Kruskal-Wallis test). **(B)** Combined density and histogram plots of relative RNA levels for each of the five Tha RABV genes color-coded by cell type and stratified by both gene identity and experimental condition. P values for differences in Neuron versus Astrocyte expression distributions are reported for each gene and experimental condition in the figure panel (Kolmogorov-Smirnov tests). (N mean %, Neurons: 26.8±0.11 sem; Astrocytes: 28.6±0.24; P mean %, Neurons: 47.3±0.08 sem; Astrocytes: 43.9±0.19; M mean %, Neurons: 22.6±0.08 sem; Astrocytes: 25.9±0.22; G mean %, Neurons: 1.19±0.02; Astrocytes: 0.79±0.03; Mean % *L* of viral RNA, Neurons: 1.9±0.01; Astrocytes: 0.99±0.02).

### Tha RABV gene expression is shaped by host cell type and viral mutations that restore host innate immune responses

We next asked whether the levels of *individual* Tha RABV genes are shaped by host cell type and 2P4M mutations, as we observed for aggregate viral mRNA expression. Interestingly, both bulk and scRNA-seq measurements of Tha mRNAs levels were not strictly explained by genome order (**Figure S3B**). *N*, *P*, and *M* were expressed at high levels and *G* and *L* at much lower levels. While cross-gene expression comparisons are potentially influenced by the methodologies used to prepare sequencing libraries, intra-gene expression comparison across scRNA profiles are unaffected by these potential technical artifacts. Therefore, we compared the relative expression of each Tha RABV gene to total viral RNAs across scRNA profiles stratified by cell type and infection condition. Because the accepted “start-stop” transcriptional model includes no role for feedback from host cell type or molecular state, we hypothesized that 2P4M mutations and host cell type should not influence the expression of each Tha gene. In contrast to our hypothesis, we observed influences of both the cell type and the Tha infection condition on relative expression of Tha RABV genes (**Figure 2B** and **Figure S3D**). Most strikingly, across all Tha RABV infection conditions, relative *L* mRNA levels were doubled in neurons compared to astrocytes. While MOI had only a marginal effect on Tha gene expression by host cell type (**Figure S3D**), *N* and *P* levels differed across Tha2P4M infected neurons and astrocytes (p < 2E-16; Mann-Whitney test), with neurons having slightly lower levels of *N* and slightly higher levels of *P* mRNAs (**Figure 2B**). These results suggest that the Tha RABV mRNA levels are sculpted by host cell type and 2P4M mutations.

### Tha RABV infection induces a cell-type-conserved pro- and anti-viral transcriptional response involving a small number of genes

Viral infections - including by various strains of RABV – can alter transcriptional regulation of the host cell genome in ways that have complex effects on host cell and tissue function^35–37^. However, systematic molecular descriptions of how RABV infection might alter the transcriptome of distinct human brain cell types – including the consequences of innate immune antagonism induced by wild-type P and M proteins – have yet to be described at single-cell resolution. Therefore, we compared genome-wide expression data across cells stratified by cell type and experimental condition to evaluate how Tha RABV infection and 2P4M-sensitive innate immune disinhibition shapes neuron and astrocyte-specific transcriptional responses.

To characterize transcriptional changes driven by Tha RABV infection and compare these changes across distinct host brain cell types, we performed genome-wide differential expression testing across cell-type-identified scRNA profiles from Tha RABV-infected (Tha(0.5)) versus “No Infection” cells, with a primary focus on neurons and astrocytes (**Figure 3** and **Figure S4A**; **Table S1; Methods**). Using false-discovery rate corrected p-values (FDR < 0.05) as a threshold, we identified 13 and 60 differentially expressed genes (DEGs) in neurons and astrocytes, respectively (**Figure 3A**). To determine the degree to which transcriptional responses to Tha RABV infection were shared or unique across neurons and astrocytes, we compared gene-specific fold-change values while also tracking gene expression levels in uninfected neurons (**Figure 3B**). Neuron and astrocyte specific responses to Tha-infection were well-correlated (r = 0.64), with a small set of human genes robustly induced in each cell type. Genes with significant reductions in astrocytes also tended to be reduced in neurons, but low neuronal baseline expression likely limited further reductions due to sensitivity of our testing. Shared, strongly induced RNAs include those from *DYNLL1*, *DYNLL2* and *HSPA8* genes and the long non-coding RNA *NRAV* (located in anti-sense orientation in the *DYNLL1* gene locus). DYNNLL1/2 are dynein-light chain subunits that interact with many other cellular proteins, bind RABV P^38–41^ and L^42^, and promote RABV transcription and replication. *HSPA8* encodes a member of the HSP70 family that is induced upon infection of RABV and other viruses, is sequestered to RABV viral factories and promotes RABV viral transcription and replication^23^. *NRAV* too has described pro-viral functions^43^. RNAs present in lower amounts in Tha RABV infected cells were largely restricted to genes encoding components of the IFN signaling, including *STAT1*, *IRF9*, *IFIT1* and *ISG15*, supporting previous observations that wild-type Tha(0.5) RABV infection condition is sufficient to suppresses IFN-induced expression of innate immune genes encoding detectors and early-stage infection effectors^20^ (**Figure S4B**). Gene expression comparisons based on ten-fold higher RABV MOI resulted in highly similar transcriptional changes (**Figure S4A**). These results demonstrate that during Tha RABV infection - in which an average of ∼3% of all mRNAs are viral (**Figure 2A**) - neurons and astrocytes tended to share changes in RNA levels from just a handful of genes: increased mRNAs encode host cell factors with previously described or candidate pro-viral functions, while reduced mRNAs encoded IFN-activated anti-viral transducer or effector proteins. In aggregate, these results suggest two functionally distinct transcriptional signatures that likely promote RABV replication in astrocytes and neurons while minimizing innate immune detection and response.

**Figure 3.**
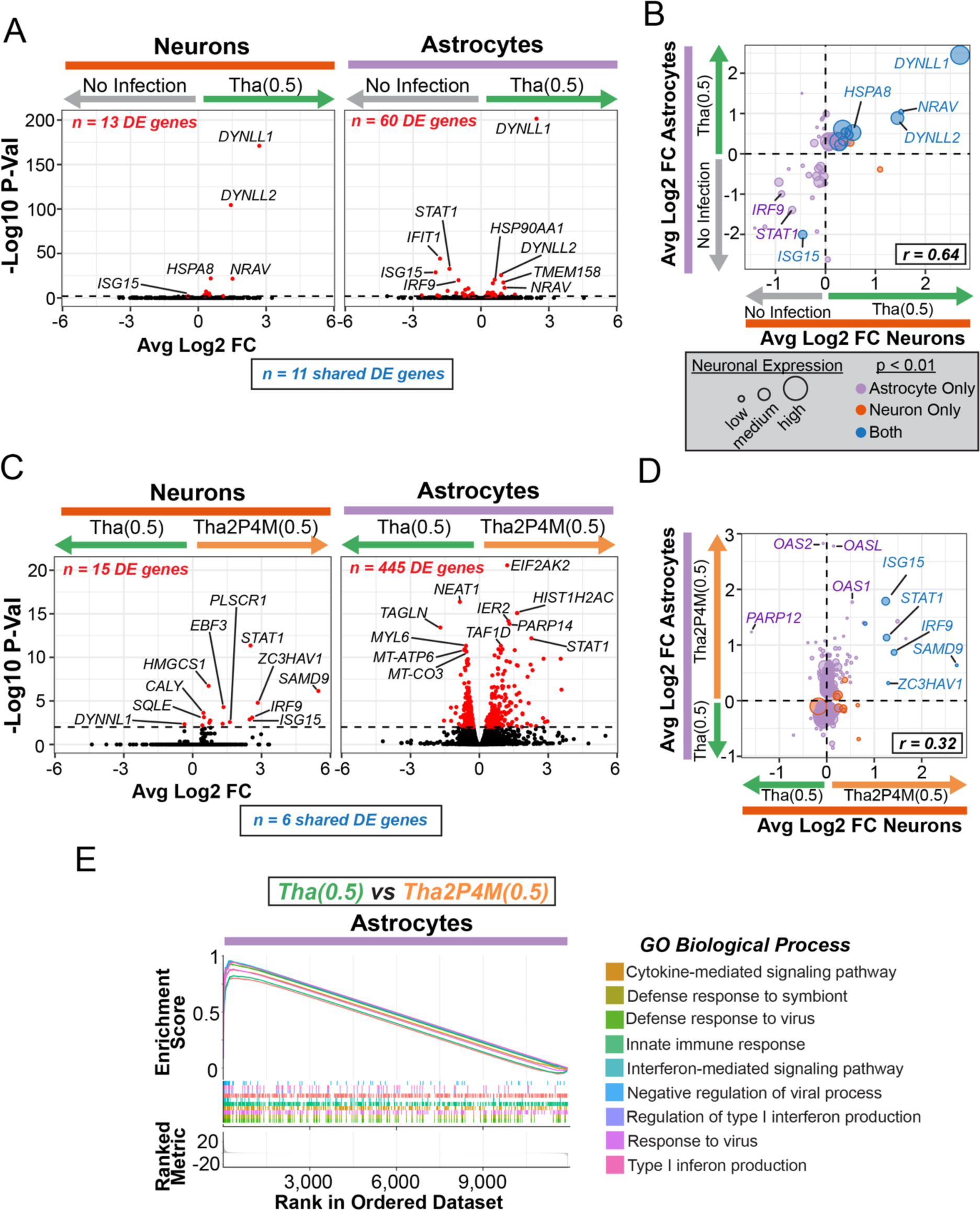
Tha2P4M mutations unmask a large-scale astrocyte-specific transcriptional response to infection. **(A-D)** Genome-wide differential expression comparisons between neurons and astrocytes across No Infection and Tha(0.5) infected (**A,B**) or infected Tha(0.5) and infected Tha2P4M(0.5) experimental conditions (**B,C**). Arrow color and direction indicates higher levels of mRNAs in that experimental condition. **(A,C)** Volcano plots of gene expression differences with select genes labelled (red, p < 0.01). **(B,D)** Scatter plots comparing log2 fold expression level changes for genes significantly altered (p < 0.01) in either neurons (orange), astrocytes (purple) or both cell types (blue). Circle size is proportional to expression level in uninfected neurons. **(E)** Gene set enrichment analysis of Biological Process Gene Ontology categories for differentially expressed genes (p < 0.01) across astrocytes infected with Tha(0.5) versus Tha2P4M(0.5) RABV. Gene set Enrichment Scores are shown for the nine most significantly enriched Biological Processes.

### Tha2P4M mutations unmask an astrocyte-specific transcriptional response to infection

To unmask and compare the transcriptional responses inhibited by wild-type Tha P and M proteins across brain cell types, we performed differential expression testing of infected, cell-type-identified scRNA profiles across Tha(0.5) and Tha2P4M(0.5) conditions (**Figure 3C** and **Figure S4A**). We observed 15 DEGs in neurons, 14 of which were upregulated (**Figure 3C**). This upregulated gene set included a subset of those same IFN-related genes that were downregulated in Tha RABV-infected astrocytes (*IRF9, ISG15, SAMD9, STAT1, PLSCR1*), suggesting this downregulation by Tha RABV infection was driven by effects downstream of wild-type P and M proteins. Astrocytes, however, exhibited significant expression changes in 445 up or down-regulated genes (**Figure 3C**). Gene-specific fold change comparisons to neurons confirmed that a small subset of the upregulated genes in Tha2P4M-infected astrocytes were cell-type conserved (n=6 of 204), while the vast majority of upregulated genes (n=198 of 204) and nearly all down-regulated genes (n=241) were unique and selective to astrocytes (**Figure 3D**). These results suggest that host brain cell type plays a profound role in the breadth and capacity for transcriptional responses triggered by innate immune signaling pathways that are typically inhibited by wild-type Tha P and M proteins. Analysis of enriched Biological Pathway gene sets in these 445 genes underscored that genes selectively induced in Tha2P4M-infected astrocytes were associated with innate immune response, viral infection, and ISGs (**Figure 3E; Methods**).

### A subpopulation of astrocytes endows a pro-viral RNA cellular environment through a massive transcriptional change

We next determined whether the large-scale transcriptional response observed in the grouped comparison of Tha2P4M infected astrocytes was driven by a specific subpopulation of astrocytes. To do this, we clustered astrocyte scRNA profiles from all experimental conditions using exclusively human genes. Our analysis identified a major (Cluster 1: n=3,759 cells; 94.6%) and a minor (Cluster 2: n=216 cells; 5.4%) astrocyte population (**Figure 4A**). The experimental composition of Cluster 2 cells strongly deviated from the full dataset, with a 3.5-fold enrichment of Tha2P4M infected astrocytes, yet all experimental conditions, including “No Infection” co-cultures, contributed cells (**Figure 4B**). Cluster 2 was also enriched for infected cells as compared to the full dataset (95% infected versus 68%; p<2.2E-16; Chi-square test)(**Figure 4B**). Based on our previous observation that a subset of astrocytes had particularly high RABV expression (**Figure 2A**), we compared viral mRNA load scores for cells in Cluster 1 and Cluster 2 (**Figure 4A**). On average, viral load scores were 6.3-fold higher in Cluster 2 than in Cluster 1 (**Figure 4C**). We conclude that a subpopulation of transcriptionally unique astrocytes supports very high levels of Tha RABV mRNA expression. Thus, for simplicity, we refer to Cluster 2 astrocytes as “pro-viral”, to connote a molecular state permissive for enhanced RABV transcription. While a small proportion of pro-viral Cluster 2 astrocytes were observed without infection (∼5%), astrocytes tended to adopt this pro-viral molecular state following Tha RABV infection, especially when the 2P4M mutations were present. Pseudotime trajectory analysis suggested that this astrocyte state transition occurs in continuous manner (**Figure S5A-C** and **Methods)**. These results imply that wild-type P and M proteins likely restrict the emergence of the pro-viral astrocyte state during Tha RABV infection.

**Figure 4.**
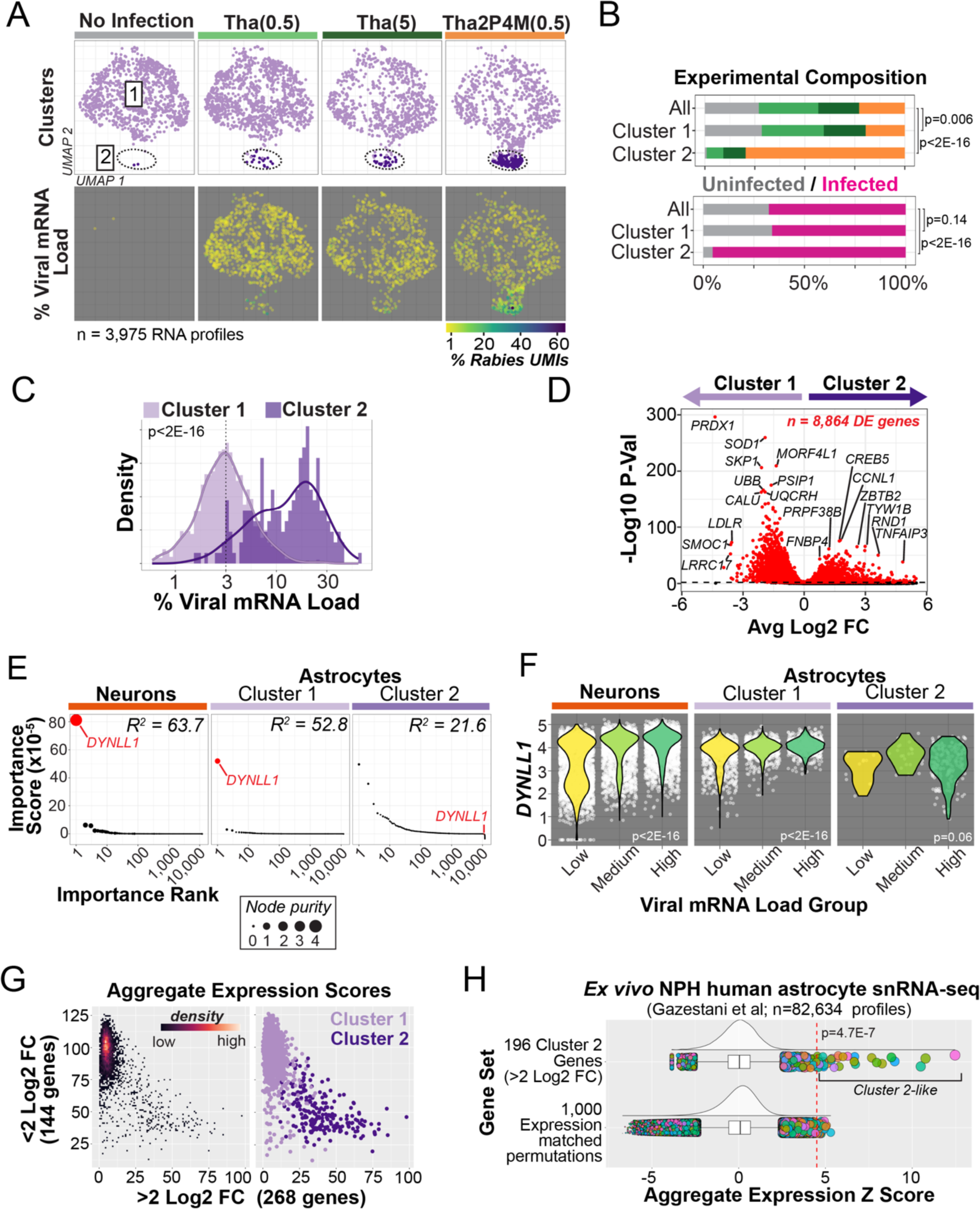
A pro-viral human astrocyte subpopulation supports high viral mRNA load and is decoupled from typical host cell transcriptional feedback through a massive alteration in RNA levels. **(A)** UMAP embedding plot of astrocyte scRNA profiles clustered exclusively by human gene expression differences and stratified by experimental condition (**Methods**). “No infection” (grey) counts include scRNA profiles from cells that belonged to the non-infected control well where RABV infection was completely absent. *Top*, color-coded cluster identities. (Cluster 1, n=3,759 cells; Cluster 2, n=216 cells). *Bottom*, heat map of viral mRNA load scores. **(B)** Bar plots comparing the composition single astrocyte RNA profiles from the full dataset versus Cluster 1 or Cluster based on experimental identity (Cluster 1 versus All: p=0.006; Chi-square test; Cluster 2 versus All: p<2.2E-16) or by infection status (Cluster 1 versus All: p=0.14; Cluster 2 versus All: p<2.2E-16). “Uninfected” counts include both uninfected “bystander” astrocytes and uninfected astrocytes from the “No Infection” condition. **(C)** Combined histogram and density plot comparing viral mRNA load distributions for Cluster 1 and 2 (Mean viral load: Cluster 1, 2.2%±0.04 sem; Cluster 2, 13.8%±0.7; p < 2.2E-16, Kolmogorov-Smirnov test). **(D)** Volcano plot of gene expression differences across Cluster 1 and 2 with select genes labelled (red, p < 0.01). **(E)** Random forest regression analysis of n=11,943 human gene RNA levels predicting viral mRNA load stratified by neurons, Cluster 1 and Cluster 2 astrocytes. Genes are rank ordered by importance score and dot size indicates node purity (**Methods**). Goodness of fit for each model is reported (R^2^). **(F)** Violin plots of *DYNLL1* expression levels stratified by both cell type and viral mRNA load groups (Neurons, p<2E-16; Astrocytes Cluster 1, p<2E-16; Astrocytes Cluster 2, p=0.05. Kruskal-Wallis tests). **(G-I)** Identification of rare “Cluster 2-like” astrocytes from single-nuclei RNA profiles derived from *ex vivo* cortical resections of living human Normal pressure hydrocephalus (NPH) surgical patients^79^. **(G)** Scatterplot of Cluster 1 or 2 astrocyte single-cell RNA profiles based on aggregate expression scores for genes up regulated in Cluster 1 (n=144 genes) or Cluster 2 (n=268 genes) (p < 0.01 and log2 fold-change > 2). *Left*, heat map of point densities. *Right*, color-coded cluster identities. **(H)** Analysis of Cluster 2 aggregate gene scores for n=82,634 single-nucleus RNA profiles of astrocytes from n=52 NPH patients (median nuclei profile per patient = 1,277). Density and boxplots show aggregate expression z-score distribution for the 196 genes shared across both datasets (*top*) versus 1000 permutations of expression-matched NPH gene sets (**Methods**). snRNA-seq profiles > 4.5 standard deviations above the mean were considered “Cluster 2 like” (n=38 profiles identified in 21 of 52 total patients; p=4.7E-7 based on permutations). In box plots, outlier nuclei profiles (> 1.5 interquartile range) are shown as dots color-coded by patient identity and dot size proportional to the z-score.

To quantify the magnitude of transcriptional changes distinguishing Cluster 1 and pro-viral Cluster 2, we performed differential expression testing. Our analysis revealed significant alteration in the RNA levels of 8,864 genes, equating to changes in ∼38% of all astrocyte expressed genes (Cluster 2 enriched: 2,343 genes; Cluster 2 depleted: 6,521 genes)(**Figure 4D**). Thus, the pro-viral state likely represents a massive change in astrocyte molecular milieu. Many genes upregulated in pro-viral Cluster 2 showed trajectory-specific expression across the astrocyte state transition in pseudotime, suggesting that the pro-viral state may be transcriptionally dynamic (**Figure S5B**). Interestingly, RNAs from at least 92 genes increased prior to pro-viral conversion (**Figure S5B**). This gene set was two-fold enriched for chromatin regulators as compared to other pro-viral genes (**Figure S5C**). Many of these chromatin regulators control cellular differentiation in other contexts, including transcription factors within the AP-1 family (*ATF3*, *FOSL1*, *FOSL2*, *JUNB* and *MAFF*^44–49^), outside the AP-1 family (*CREB5*, *CSRNP1*, *EGR2*, *ETS1*, *FOXS1*, *HEY2*, *ISL2*, *KLF5*, *MEF2C*, *NFATC2*, *PBX4*, *TEAD4*^50–76^), the histone variant *CENPA*^77^ and the histone modifier *JADE1*^78^.

To determine whether high viral mRNA load was more likely the cause or the consequence of pro-viral astrocyte transcriptional state, we performed two complementary analyses. First, we inspected the relationship of pro-viral identities, pro-viral gene expression and viral mRNA load across pseudotime (**Figure S5B**). Acquisition of pro-viral identities and up-regulation of many pro-viral RNAs tended to precede viral mRNA load increases. Second, we repeated differential expression testing after downsampling scRNA profiles such that viral mRNA load distributions were equivalent across clusters (**Figure S5D**; **Methods).** Gene-specific fold change values were highly correlated across the full and downsampled analysis (r = 0.82), with 32% of significant genes shared in both comparisons (**Figure S5E,F**). The vast majority of other genes passing the significance threshold in the full dataset but had similar, non-significant directional changes in the downsampled data (**Figure S5F**), likely due to the drop in test sensitivity from far fewer cells in the analysis (n= 276 versus 3,975 scRNA profiles). These analyses suggest that high viral mRNA load is more likely a consequence rather than cause of the pro-viral state.

We next tested whether the pro-viral state affects the transcriptional coupling between RABV and host cell genome during infection. To do this, we performed a random forest-based regression analysis, using viral mRNA load scores as the dependent variable and 11,943 human genes as independent variables (**Methods**). In neurons and Cluster 1 astrocytes, regression models explained high levels of variance in viral mRNA load (R^2^ = 63.7 or 52.8) and both analyses highlighted a privileged role of *DYNLL1* RNA levels in predicting viral mRNA load (**Figure 4E**). In Cluster 2 astrocytes, the regression model explained far less variance (R^2^ = 21.6) and *DYNLL1* levels were no longer predictive. Models run without *DYNLL1* explained far less variance in viral mRNA levels for neurons and Cluster 1 astrocytes (R^2^ = 39.3 or 32.6) but not for Cluster 2 astrocytes (R^2^ = 20.6). Indeed, stratification of scRNA profiles into low, medium and high viral mRNA groups illustrated that *DYNLL1* RNA levels grew with viral mRNA load scores in neurons and Cluster 1 astrocytes but not in pro-viral Cluster 2 astrocytes (**Figure 4F**). These results suggest that the molecular properties of pro-viral Cluster 2 astrocytes fundamentally alter the transcriptional coupling between RABV and host cell genome, including the typical strong association between *DYNLL1* RNA and RABV transcriptional output.

### Identification of pro-viral Cluster 2-like astrocytes in the human brain

The presence of ∼5% of uninfected astrocytes in Cluster 2 (including eight “bystander” uninfected astrocytes and two astrocytes from the “No Infection” condition), hinted that this pro-viral astrocytic transcriptional state might exist independent of RABV infection and motivated us to explore the possibility that Cluster 2-like astrocytes might be present in human brains. To evaluate this possibility, we took advantage of a snRNA-seq dataset acquired from flash-frozen cortical tissue resected during Normal Pressure Hydrocephalus (NPH) surgery^79^, enabling analysis of *ex vivo* astrocyte RNA profiles from 52 living patients. Therefore, these cortical samples were collected without the confound of post-mortem interval nor acute central nervous system infection. To search for Cluster 2-like astrocytes in the 82,634 NPH snRNA-profiles from astrocytes, we developed an aggregate expression score approach, leveraging genes highly enriched (> 2 log2 fold-change) in pro-viral Cluster 2 astrocytes to create a “discovery” gene set (**Figure 4G,H**; **Methods**). As expected, aggregate expression of this Cluster 2 “discovery” gene set (n=268 genes) clearly distinguished Cluster 1 and Cluster 2 scRNA profiles in a manner independent from our original analysis, confirming the approach’s sensitivity (**Figure 4G**). We then calculated aggregate expression scores for NPH astrocyte snRNA-seq profiles based on the 196 “discovery” genes expressed in both datasets. To contextualize the distribution of “discovery” gene set scores and define a false-discovery threshold, we performed a similar analysis on n=1,000 permuted gene sets of 196 genes matched for expression level. We then calculated gene set-specific z-scores to directly compare the “discovery” and permutation results (**Figure 4H**; **Methods**). In support of Cluster 2-like astrocytes present in living human cortex, we observed a small number of snRNA profiles with very high “discovery” set z scores (up to 12.5 standard deviations above the mean) not observed in the permutations. Using a z-score threshold of 4.5 (corresponding to a p value cut off of 4.7E-7 based on the permutations), we identified n=38 Cluster 2-like astrocytes contributed by 21 of 52 total patients (n=1 to 4 Cluster 2-like NPH astrocytes per patient, ranging from 7 to 4,990 total astrocyte snRNA profiles per patients (median: 1,278). We reasoned that if these 38 single-nucleus RNA profiles were indeed Cluster 2-like, they should also be enriched for an independent gene set with enriched expression in pro-viral Cluster 2 astrocytes. Indeed, a “validation” gene set of 521 genes with RNAs 1-2 log2 fold-enriched in Cluster 2 astrocytes also had enriched expression in the 38 snRNA-profiles as compared to all other NPH astrocytes (n= 82,596: p < 2E-16, chi-square test). The degree of “discovery” and “validation” gene set enrichment in NPH astrocytes corresponded to the predictions from pro-viral Cluster 2 astrocytes (NPH expression enrichment: “discovery”, 2.8; “validation”, 1.2)(**Figure S5G**). Collectively, these analyses provide preliminary support for a rare (∼1 in 2,200) population of pro-viral Cluster 2-like astrocytes in human cortex without acute infection of the central nervous system (**Figure S6A,B**).

### Transcriptional analysis identifies major cell-autonomous and nonautonomous functional changes of pro-viral astrocytes

We next sought to analyze how the massive transcriptional changes distinguishing Cluster 1 from pro-viral Cluster 2 astrocytes might alter both intercellular interactions with other cell types and cell-autonomous astrocyte function. Importantly, we found no evidence that the pro-viral transcriptional state we observed in Tha2P4M infected human astrocytes was previously documented in cultured or *ex vivo* rodent and human astrocytes experimentally driven into “reactive” states through interferon^80,81^ or lipopolysaccharide exposure (**Figure S7A**).

To understand to what degree the pro-viral transcriptional state affects astrocyte signaling with other co-cultured brain cell types, we leveraged CellPhoneDB(v5.0), a database of pairwise-protein interactions that includes both secreted and extracellular ligands in relationship to receptor complexes. We focused our analysis on genes with > 1 log2 fold-enrichment in either Cluster 1 (n=2,186) or Cluster 2 (n=877). CellPhoneDB uses cell-type-assigned scRNA-seq data and a gene-shuffling permutations to identify “relevant” ligand-receptor interactions that are selectively enriched in target gene sets over background expression in other cell types (p < 0.05 based on permutations). CellPhoneDB analysis revealed that pro-viral Cluster 2 astrocytes tended to express fewer “relevant” ligand and receptor genes than Cluster 1 astrocytes, suggesting a global reduction in the capacity for intercellular signaling (**Figure 5A**). Interestingly, the more minor subset of ligands upregulated in pro-viral astrocytes include immune effectors that function as secreted chemokines (*IL34*, *LIF*, *EDN1*) or extracellular glycoproteins (*VCAM1* and I*CAM1*). In some cases, ligands or receptors from the same signaling family exhibited bi-directional changes in RNA levels. For example, *WNT3A* and *VEGFA* (ligands) and *EPHA4* (receptor) were enriched while *WNT7B* and *VEGFB* (ligands) and *EPHB1* (receptor) were reduced in pro-viral Cluster 2 Astrocytes. Analysis of “relevant” ligands and receptors in the context of RNAs expressed by other co-cultured brain cell types suggested that pro-viral Cluster 2 astrocytes (as compared to Cluster 1 astrocytes) exhibit reduced signaling capacity to all major cell types to approximately the same degree. These data also imply that pro-viral Cluster 2 astrocytes are more “cut-off” from receiving interactions from other cell types (save from other pro-viral and Cluster 1 astrocytes)(**Figure 5B**). These inferences based on RNA levels may suggest that pro-viral Cluster 2 astrocytes are poised to send and receive fewer total cellular interactions yet highlight a small number of key pathways through which astrocytes in the pro-viral molecular state might signal to other brain cell types.

**Figure 5.**
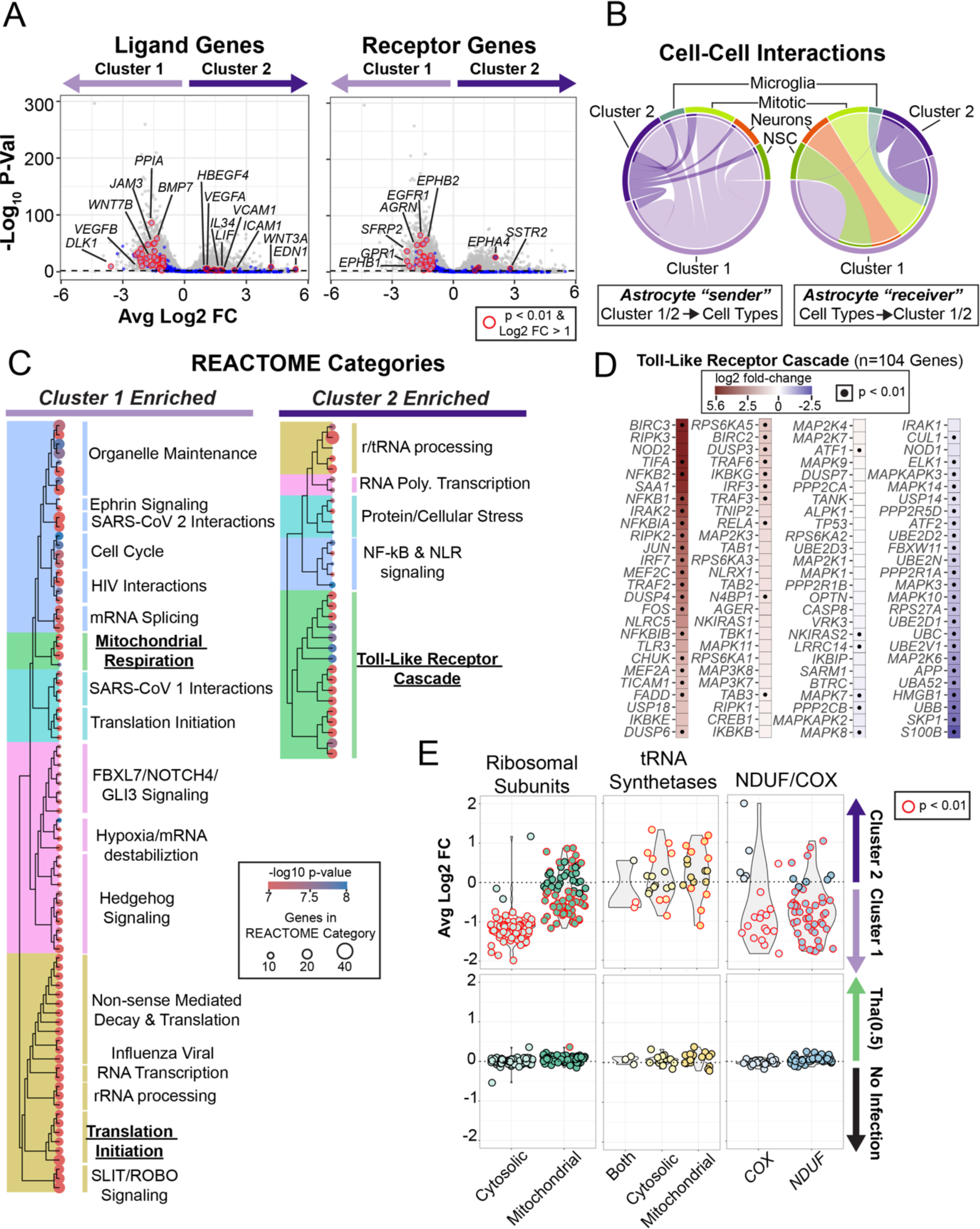
Gene expression changes in pro-viral human astrocytes imply reduced inter-cellular interactions, activation of toll-like receptors and complex changes in protein translation and mitochondrial respiration. **(A,B)** Gene expression inference of alterations to inter-cell type interactions between Cluster 1 and Cluster 2 astrocytes using CellPhoneDB (CPDB) v5.0^105^, a database of 1,419 molecular interactors (**Methods**). **(A)** Volcano plot-based visualizations of ligand (*left*) or receptor (*right*) gene expression changes between Cluster 1 and Cluster 2. Ligand or receptor genes significantly enriched in either cluster (p<0.01) and significantly “relevant” (after CPDB permutation; p < 0.05) are circled in red (Ligand genes: Cluster 1, n=46; Cluster 2, n=11; Receptor genes: Cluster 1, n=32; Cluster 2, n=5). Other ligand or receptor genes are shown in blue and all other genes in gray. **(B)** Chord diagrams showing the number of CPDB relevant interactions from (*left*) or to (*right*) Cluster 1 or Cluster 2 astrocytes for other cell types. The proportional size and color of the outer ring corresponds to the relative number of interactions by cell type. The color and proportion of the inner ring corresponds to interactions from or to Cluster 1 or Cluster 2 astrocytes. **(C)** Tree plot clustering of enriched REACTOME gene categories (Cluster 1: p < 0.000001, n=84 categories; Cluster 2: p < 0.01, n=32 categories) based on genes upregulated in Cluster 1 (*left;* n=6,551) or Cluster 2 (*right;* n=2,347)(p<0.01). Hierarchical clustering results reflect shared genes driving each REACTOME category association. Dot size scales with the gene number in each category. Heat map indicates p value of each included category. Underlined REACTOME categories are the focus of panels (**D**) and (**E**). **(D)** Heat map plot reflecting log2 fold-changes in Cluster 2 of each gene in the Toll-Like Receptor Cascade REACTOME category (R-HSA-168898; n=104 genes). Dots indicate significant expression changes (p < 0.01). **(E)** Combined violin and dot plots reflecting gene-specific log2 fold-changes for gene sets encoding proteins related to protein translation (*left* and *middle*) or mitochondrial respiratory complexes I and IV (*right*). *Top row*, Cluster 1 versus Cluster 2 astrocytes. *Bottom row,* No Infection versus Tha(0.5) RABV infected astrocytes. Ribosomal subunit genes are stratified by whether the encoded protein is a component of cytosolic (n=85 genes) or mitochondrial (n=78 genes) ribosomes. tRNA synthetase genes are stratified by whether the encoded enzyme aminoacyl-tRNAs used for translation in the cytosol (n=17 genes), mitochondria (n=17 genes) or both ribosome types (n=3 genes). Nuclear gene components or assembly factors of the cytochrome c oxidase (*COX*) family (n=23 genes), Complex IV. The NADH:ubiquinone oxidoreductase (*NDUF*) family (n=47 genes) encoding Complex I. Red outlines indicate significant expression changes (p <0.01).

We next leveraged gene expression changes to build testable hypotheses for how the pro-viral transcriptional state might affect cell-autonomous function of human astrocytes. Pro-viral Cluster 2 astrocytes expressed higher levels of annotated human innate immunity genes^82^ (n=837) including those stimulated by IFN (n=244) (**Figure S7B,C**). To find additional themed structure in the ∼38% of astrocyte-expressed genes differentially expressed across Cluster 2 and Cluster 1 astrocytes, we performed hierarchical clustering of REACTOME categories based on shared genes up regulated in Cluster 1 (n=6,521 genes; n=84 categories, p <1E-6) or Cluster 2 (Cluster 2, 2,343 genes; n=32 categories, p<1E-3)(**Figure 5C**). These results implicate a diverse set of cellular processes affected by the large-scale restructuring of the pro-viral astrocyte transcriptome. Due to the number and complexity of these changes, we focused further analysis on the following salient gene sets to more fully understand how astrocytes in particular respond to infection: 1) genes controlled by “Toll-like receptor cascades” (**Figure 5D**) and genes encoding machinery for 2) protein translation and 3) mitochondrial respiration (**Figure 5E**). (“Toll-like receptor (TLR) cascades” is likely a proxy for gene sets activated by the IRF3/IRF7/NF-κB transcription factor complex, which in the case of RABV infection, is primarily driven through activation of viral sensors such as TLRs and RIG-I-like receptors (RLRs)^83,84^).

To determine how the molecular milieu of Cluster 2 astrocytes is sculpted by RNA level changes of genes controlled by “Toll-like receptor cascades”, we ranked fold-changes for 108 associated genes (**Figure 5D**). This analysis highlighted the striking role for transcriptional induction of both NF-κB and immediate early gene (IEG) transcription factors and regulators (NF-κB: *NFKB1*, *NFKB2*, *NFKBIA*, *NFKBIB*; IEG: *JUN*, *FOS*) as well as induction of gene targets controlled by both NF-κB and IEG transcriptional complexes (NF-κB: *BIRC3*, *BIRC2*, *NOD2*, *RIPK3*, *IRAK1, TRAF2;* IEG: *DUSP4, DUSP6, DUSP3*). RNAs from this group with lower relative levels in the pro-viral state encode MAP kinases and regulators (*MAP2K6*, *MAPK10*, *MAPK3*, *MAPK14*, *MAPKAPK3*, *MAPK8*, *MAPK7*) as well as ubiquitin and ubiquitin regulators (*UBB*, *UBA52*, *UBE2V1*, *UBC*, *UBE2D1*, *UBE2N*, *UBE2D2*). Thus, viral sensor activation may contribute to the pro-viral state through relative increases of NF-κB genes and targets and relative reductions of RNAs that might alter MAPK kinase activity and ubiquitin-based protein degradation.

To determine how transcriptional alterations of genes encoding protein translation and mitochondrial function might contribute to the function of pro-viral astrocytes, we assessed expression-level changes of ribosomal subunits, tRNA synthetases and the COX and NDUF gene families encoding Complex IV and Complex I of the electron transport chain, respectively (**Figure 5E**). Pro-viral astrocytes exhibit potent reductions in RNAs encoding nearly all subunits of the cytosolic ribosome (n=85 genes), and bi-directional regulation of mitochondrial ribosome subunits (n=78) and tRNA synthetases (n=37 genes). Most members of the COX and NDUF gene families had reduced expression in pro-viral astrocytes. These transcriptional changes were highly-selective for pro-viral astrocytes, as RNA levels associated with these genes were not different across No Infection and Tha(0.5) RABV infected astrocytes (**Figure 5E**). These results suggest that pro-viral astrocytes likely have severely reduced or dysregulated capacity for protein translation and oxidative phosphorylation, potentially indicating a mechanism of “host shut off” based on RNA level changes. Tha RABV infection with wild-type P and M protein efficiently inhibit these changes.

### Microglia indirectly respond to infection of other cell types through interferon-stimulated genes potentiated by 2P4M mutations

Lastly, we explored how RABV infection-resistant microglia indirectly respond to infection properties of other susceptible co-cultured brain cell types and how these transcriptional responses associate with Tha RABV titer or innate immune signaling disinhibited by 2P4M mutations. To do this, we re-clustered microglia scRNA-seq profiles and tracked the experimental origins of each library (**Figure 6A**). Three distinct analysis or visualization approaches – including UMAP embeddings (**Figure 6A** and **Figure S8A**), principal components analysis of shared expression variance (**Figure 6B** and **Figure S8B**) and differential expression analysis (**Figure S8C**) – revealed that exposure to Tha RABV particles and the milieu of neighbouring infected cells affect a graded transcriptional response of interferon-induced genes in microglia (e.g. *SNX10*, *GBP1*, *IFIT2*, *ACSL1*)(**Figure 6B**). The number and degree of interferon-induced transcriptional differences were largest in the Tha(5) condition, followed by Tha2P4M(0.5) and then Tha(0.5)(**Figure 6B-D** and **Figure S8C**), as evidenced by example genes (*ISG15, IDO1, and IFIT3*)(**Figure 6E**). Consistent with rapid induction and degradation^85^, RNAs encoding interferon proteins were not detected 48 hours after viral infection, obfuscating the cellular source of interferons in our analysis. Importantly, in titer-matched experiments, the interferon-related transcriptional response in microglia was larger with 2P4M mutations (**Figure 6C**), suggesting that disinhibited innate immune signaling and/or higher levels of viral RNA in other cell types potentiate an interferon-stimulated gene responses in microglia. However, analysis of cytokine gene RNA levels in neurons and astrocytes found little support for this model (**Figure S8D**). The majority of cytokine ligand and receptor RNAs were expressed at high levels in microglia themselves (**Figure S8E**). In many cases, both ligands (e.g. *CCL3*, *CCL4*, *CCL5*, *CCL8* and *IL1B*) and receptors (e.g. *CCR5*, *CCR1*, *IFNGR1* and *TNFRSF1A*) were expressed across experimental conditions in a manner similar to ISGs (**Figure S8F**).

**Figure 6.**
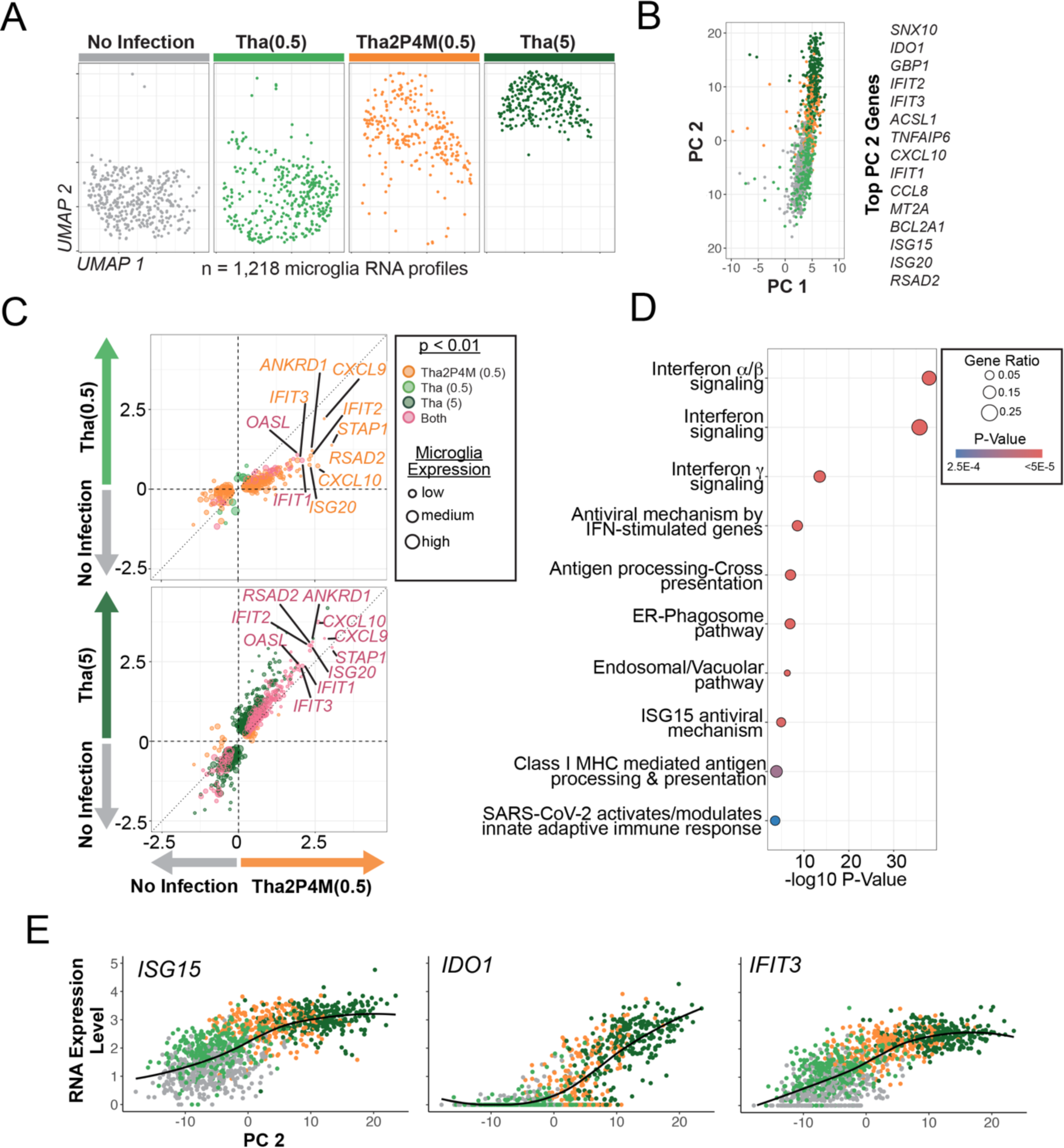
Uninfected microglia respond to Tha RABV infection through graded interferon stimulated gene transcription potentiated by higher viral titer and disinhibited innate immune signalling of other infected cell types. **(A)** UMAP embedding plot of microglia scRNA profiles stratified by experimental condition (**Methods**)(n=1,218 cells). **(B)** Principal component analysis of variably expressed microglia genes. *Left*, cell component loadings of PC 1 (contamination of neuronal RNAs; **Figure S8B**) versus PC 2 (interferon stimulated gene expression; **Figure S8B**). *Right*, n=15 genes with the highest gene loadings for PC 2. **(C).** Scatter plots comparing log2 fold expression level changes of differentially expressed genes (p < 0.01) across No Infection microglia and microglia from various experimental conditions (*top*, Tha(0.5) versus Tha2P4M(0.5); *bottom*, Tha(5) versus Tha2P4M(0.5)). Color reflects significant genes shared across comparisons (pink) versus those unique to experimental conditions. The dotted unity line (slope = 1, y intercept = 0) serves as a reference to indicate equivalent log2 fold changes across conditions. **(D)** REACTOME pathway analysis plot based on n=202 shared genes found differentially expressed in comparisons of both Tha2P4M(0.5) and Tha(5) to No Infection microglia. **(E)** Normalized gene expression plots for three representative interferon stimulated genes plotted against principal component 2 scores and color-coded by experimental condition.

## Discussion

In this work, we characterized how the innate immune signaling inhibition conferred by RABV P and M proteins shapes the transcriptional properties of the host and viral genomes across human neurons and glia. While molecular mechanistic analyses in cell lines have described the potent and synergistic ways that RABV M and P proteins inhibit the induction and response to IFNs^17–19,28–31^, the consequences of innate immune inhibition or disinhibition on host-virus relationships were unknown. Here we used an experimental and analytical framework with several key attributes to characterize these consequences in major human CNS cell types, revealing important new insights into RABV-host cell interactions and how innate immune antagonism shapes viral infection properties. These attributes include 1) the use of Tha RABV derived from a field isolate (that exhibits profound innate immune inhibition as compared to attenuated RABV strains more adapted to cell culture) and 2P4M mutations (that selectively and potently strip away major RABV innate immune inhibitory mechanisms); 2) co-cultured human stem-cell derived brain cell types, enabling human cell biology to be studied in a relevant model of cross-cell type signaling; 3) single-cell RNA-seq phenotyping in which mRNA level across the RABV and human genomes are measured at cellular resolution; and 4) novel analytical methods to qualitatively group infected and uninfected scRNA-seq profiles across heterogenous infection states and the use of “viral mRNA load” as a quantitative proxy for infection state progression. (Viral mRNA load varied considerably in infected cells after 48 hours, from 0.4 to 65% viral mRNA, highlighting the importance of tracking viral mRNA load when experimentally phenotyping individual cells). Viral mRNA load distributions suggested no evidence of secondary RABV infections, consistent with the slower replication rates associated with isolate RABV and primary brain cells.

Our results suggest two ways in which Tha RABV gene expression is not anticipated by the accepted “start stop” transcriptional model. First, in both bulk mRNA and scRNA-seq based datasets, Tha RABV mRNA levels were not strictly explained by genome order, with *N*, *P* and *M* levels high (∼24-46% of viral mRNAs) and *G* and *L* levels much lower (∼1-2%). While mRNA capture and library preparation methods can complicate cross-gene RNA level comparisons, the lack of strict genome-ordered expression across independent methods suggests other factors (such as post-transcriptional regulation) may need to be considered to fully understand variable levels of RABV transcripts in infected cells. Second, scRNA-seq analysis suggests a clear if subtle role for host cell type in shaping RABV gene expression. Most saliently, *L* mRNAs were observed on average at double the levels in neurons as compared to astrocytes, differences that could be driven by either molecular or physiological differences in host cells. Future experiments are warranted to fully validate and understand the functional consequences of cell-type-specific RABV gene expression differences and to identify the mechanisms driving these effects.

In Tha RABV-infected neurons and astrocytes, we discovered largely conserved bi-directional transcriptional programs involving ∼60 human genes that may promote infection progression by both boosting RNAs encoding known pro-viral proteins or reducing RNAs encoding anti-viral transduces or effectors. For example, of the four genes robustly induced by Tha RABV infection (*DYNLL1, DYNLL2, NRAV and HSPA8;* **Figure 3A**), all have been shown to promote viral replication in some contexts^42^. Importantly, the products of the protein-coding genes (DYNLL1, DYNLL2 and HSPA8) are bound and/or sequestered by RABV proteins during infection. DYNLL1/DYNLL2 were originally described as light-chain dynein subunits but are now appreciated as “hub” proteins that perform myriad cellular roles through hundreds of unique binding partners^86,87^. DYNLL1/DYNLL2 are targeted by RABV P^38–41^ and L^42^ as well as proteins from diverse other viruses^88^. The long non-coding RNA *NRAV* spans exons 2 and 3 of *DYNLL1* gene locus in the anti-sense orientation on chromosome 12 and promotes influenzae replication and virulence^43^. The functional effects of *NRAV* on RABV pathogenesis are yet to be determined. *HSPA8* encodes a member of the HSP70 family, a host cell protein recruited to RABV Negri Bodies (the specialized viral organelles in which viral transcriptional and replication occur) and promotes RABV replication^89^. Our data thus suggest that RABV (and perhaps other viruses) evolved to sequester host cell factors transcriptionally regulated by hijackable negative feedback systems, enabling pro-viral host factor production to match viral demand as infection proceeds. Indeed, our analysis demonstrated that, compared to thousands of other human genes, *DYNLL1* mRNA levels uniquely scale with viral mRNA load in neurons and Cluster 1 astrocytes (**Figure 4E,F**). Previous work has identified a potential mechanism: DYNLL1 sequestration by P and L might disinhibit DYNLL1-based negative regulation of the ASCIZ transcription factor which promotes *DYNLL1* transcription^90^. More experiments are required to understand these transcriptional feedback systems, including the mechanism through which *HSPA8* RNA levels are increased following RABV infection. The genes potently downregulated by Tha infection include RNAs encoding proteins with central roles in anti-viral responses. These include anti-viral effectors IFIT1 and ISG15, as well as core components of the ISGF3 transcriptional complex (STAT1 and IRF9) responsible for inducing interferon expression^91^. These results offer insight into the mechanisms through which Tha RABV infection completely blocks interferon-stimulated gene expression, even after exogenous IFNα stimulation^20^. While more work is necessary to determine exactly how anti-viral RNAs are reduced during Tha RABV infection, wild-type P and M proteins are likely involved, as an overlapping anti-viral gene set is strikingly induced in neurons infected by Tha2P4M (**Figure 3C**).

One major observation of our study is that 2P4M-based disinhibition of innate immunity results in a mean doubling of viral mRNA load across infected cell types (**Figure 2A** and **Figure S3C**). Three pieces of evidence suggest that increased viral transcription is an indirect effect of the disinhibited innate immune milieu rather than a direct effect of the 2P4M mutations on the efficacy of the viral transcription complex involving P protein and the RNA-dependent RNA polymerase L. First, the 2P4M mutations sit at residues independent and distant from those known to bind or modulate L function^15,20,92^ (**Figure S1D-F**). Second, Tha and Tha2P4M have indistinguishable replication properties in BSR-T7 cells that lack an IFN response^20^. Third, P/M-based restriction of RABV transcription is cell type and state-specific, as observed in the “pro-viral” astrocyte subpopulation, which had on average six-fold higher viral mRNA load scores than other neurons and astrocytes (Cluster 2; **Figure 4A,C**). Therefore, one important consequence of RABV innate immune inhibition is likely to restrict viral transcription.

Our discovery and characterization of “pro-viral” human astrocytes deepens the growing appreciation of how astrocytes contribute immune functions in the human CNS^11,93,94^ including as viral reservoirs^95–98^. Our *in vitro* analyses suggest that the “pro-viral” molecular state is transcriptionally dynamic and collectively involves ∼38% of astrocyte expressed genes. Four lines of evidence suggest that the “pro-viral” state can exist independently of RABV infection and that this state is more likely the cause rather than the consequence of high viral mRNA load: 1) The presence of 5% uninfected “pro-viral” astrocytes (including those from uninfected culture wells); 2) Changes in the levels of many “pro-viral” genes that precede the transition into “pro-viral” Cluster 2 identities and or do not scale with viral mRNA load. These “early” genes include transcription factors and other chromatin modifiers known to control cellular differentiation in other systems; 3) the observation that “pro-viral” gene expression signatures remain after controlling for viral load differences; and 4) the presence of “pro-viral”-like single-nucleus RNA profiles sampled from cortical biopsies of presumably RABV free NPH surgical patients (∼1 out of 2,200; observed in 21 of 52 patient samples). Importantly, Tha RABV infection marginally increases the probability of transition into the pro-viral state and that probability is dramatically increased when the 2P4M mutations are present (**Figure 4A,B; Figure S5B; Figure S6A,B**). These data may suggest that inhibition of pro-viral astrocyte state emergence during infection is an important feature of RABV innate immune inhibition in the human CNS.

We leveraged gene expression data to draw a preliminary description of how the large-scale “pro-viral” state might affect astrocyte function and cellular interactions. Our analysis suggests that the “pro-viral” state tends to reduce the number of molecular opportunities for cell signaling to other brain cell types and the only unique interactions “pro-viral” astrocytes are poised to receive are from other astrocytes (in contrast to other infected astrocytes, which maintain receptors to interact with all other cell types in our co-cultures). Our analysis highlights a number of key signaling molecules including both ligands (*HBEGF4*, *VEGF4*, *IL34*, *VCAM1*, *ICAM1*, *WNT3A* and *EDN1*) and receptors (*EPHA4*, *SSTR2*) that merit further investigation for “pro-viral” astrocyte function^99^. Cell autonomously, “pro-viral” gene expression changes indicate a dramatic increase in ISG and NF-κB target gene expression, predicted to alter MAPK function and protein ubiquitination, among other changes. We also observed cellular mRNA level changes consistent with potent reductions in ribosomal machinery for protein translation and mitochondrial machinery for oxidative phosphorylation. The changes could reflect an astrocyte-specific form of “host shut off”. (To our knowledge, “host shut off” molecular programs have not been previously described either in astrocytes or for RABV infection of any cell type). The uncoupled transcriptional feedback between *DYNLL1* levels and viral mRNA observed in “pro-viral” astrocytes (but not in other cell types) may be an important feature of this process (**Figure 4E,F**). Astrocyte-based abortive infections occur at higher frequency in lab-attenuated versus field isolate RABVs^100^. Thus, it will be important to explore whether the “pro-viral” state we observed at higher rates following Tha2P4M infection might represent a transcriptional snapshot of the abortive infection process^58^. High levels of viral mRNA and with reduced protein translation could in theory help “pro-viral” astrocytes create an immune-stimulatory environment through viral RNA while minimizing the generation of infective virions. More work – including through astrocyte infection fate-mapping experiments – is necessary to clarify the functions of key pro-viral state genes and whether the pro-viral state supports abortive infections or other outcomes^101^.

Our study also sheds a new light on the role of microglia. Our analysis suggests that microglia are almost exclusively recalcitrant to RABV infection (∼1% infected, n = 10 of 859; **Figure 1G; Figure S8A**). The susceptibility of human microglia to RABV infection has been a topic of scientific debate^37,102,103^. Our findings indicate that human microglia do not play major roles in the replication and spread of dog-adapted RABV (the major cause of RABV infection in humans). The presence of viral mRNAs in the 1% of microglia we classified as infected may also be attributed phagocytosed material from infected cells. Further, the degree of microglial interferon-induced transcriptional differences was largest in the Tha RABV(5) condition, followed by Tha2P4M(0.5) and then Tha RABV(0.5)(**Figure 6** and **Figure S8C**) a signature that could not be simply explained by cytokine expression in neurons and astrocytes (**Figure S8D-F**). These results may suggest that graded microglia ISG and cytokine responses may be controlled through sensing of RABV pathogen-associated or danger-associated molecular patterns (PAMPs and DAMPs). These data are consistent with model in which microglia ISG and cytokine RNAs are primarily driven by transient interferon release by other brain cell types or direct microglia sensing of RABV-associated molecular patterns without direct infection. The larger ISG responses microglia in the Tha2P4M(0.5) versus Tha(0.5) conditions could reflect 2P4M-dependent increases in free RABV particles (a potential source of PAMPs/DAMPs) in media or signaling responses downstream of 2P4M infections in other cell types. Future experiments should directly test these hypotheses including the identification of possible signaling pathways responsible for RABV recognition by microglia.

Our study has important limitations and follow-up studies are necessary to more fully understand the biological results we reveal. For example, our genome-wide molecular phenotyping was restricted to polyadenylated RNA levels. To more fully understand the molecular consequences of isolate RABV infection and “unmasked” innate immune signaling, single-cell measurements of RABV negative-stranded RNA genomes, full-length host and viral RNA isoforms, and non-coding RNAs (including RABV leader and trailer RNAs) will need to be measured. While the 2P4M mutations profoundly disinhibit innate immune signaling^20^, this disinhibition may be incomplete. For example, the serine at residue 179 of P has been shown to be necessary for sequestering IKKε kinase - a key regulator that controls interferon transcription by activating the IRF3/7 transcription factors – and this serine is present in both wild-type P and the 2P mutant. Additionally, more work is required to fully understand the function of “pro-viral” astrocytes and the roles contributed by specific genes induced in the pro-viral state. Future experiments should 1) locate and quantify “pro-viral” astrocytes through molecular anatomical analysis of human brain tissue; 2) characterize the temporal progression and fate of the “pro-viral” state across different conditions (infection both other viruses, aging and neurodegeneration, for example) and 3) functionally assess how the “pro-viral” state affects astrocyte physiology, cell signaling and nuclear chromatin organization. Finally, it is unknown if pro-viral astrocytes are present in other species. Discovery of a similar astrocyte state in mouse, for example, would present a powerful opportunity for functional analysis *in vivo*.

Another limitation stems from the sex differences in the karyotypes of the cell types used in this study. The iPSC-derived microglia used had a male karyotype, whereas the neurons and astrocytes derived from human neural stem cells (hNSCs) had a female karyotype^104^. These sex-specific differences may influence cellular responses and contribute to some of the observed outcomes. Future studies would benefit from a systematic examination of how sex-linked genetic and epigenetic differences and genetic variation affect RABV infection dynamics in human brain cell models.

Despite these limitations, our single-cell RNA-seq analysis using the Tha/Tha2P4M RABV study system in human brain cell-type co-cultures provides a powerful approach to discover key features of host-RABV interactions and a road map for building testable hypotheses regarding the dynamics and key molecular interactions that drive neurotropic infection in the brain.

## Acknowledgements

We thank Anne Danckaert and Nathalie Aulner (UTechS Photonic BioImaging (PBI) core at Institute Pasteur Paris) for help with the image acquisition using the Opera Phenix Plus High Content Screening System (Perkin Elmer) and data analysis using the Columbus software (Perkin Elmer). PBI is a member of the national infrastructure France-BioImaging supported by the French National Research Agency (ANR-10-INBS-04). We acknowledge financial support of the Région Ile-de France (program DIM1Health, to H.B.), the Institut Pasteur (H.B. and L.F.), the Sloan Foundation (to A.S.) and the NIH Brain Initiative (R01 MH130464, to A.S.). We thank all other members of the Saunders and Bourhy’s labs for helpful discussions of this work.

## Author contributions

Conceptualization (LF, KY, HB, AS); Data Curation (ZG, JPC); Formal Analysis (KY, GC, LBA, AS); Funding acquisition (HB, AS); Investigation (LF, TK, LK); Methodology (LF, KY, AS); Project Administration (HB, AS); Resources (LF, VL, FG, MH); Software (ZG, AS); Supervision (HB, AS); Validation (LF, AS); Visualization (KY, GC, AS); Writing – original draft (LF, AS); Writing – review & editing (LF, KY, GC, HB, AS).

## Supplemental Figures

**Supplemental Figure 1.**
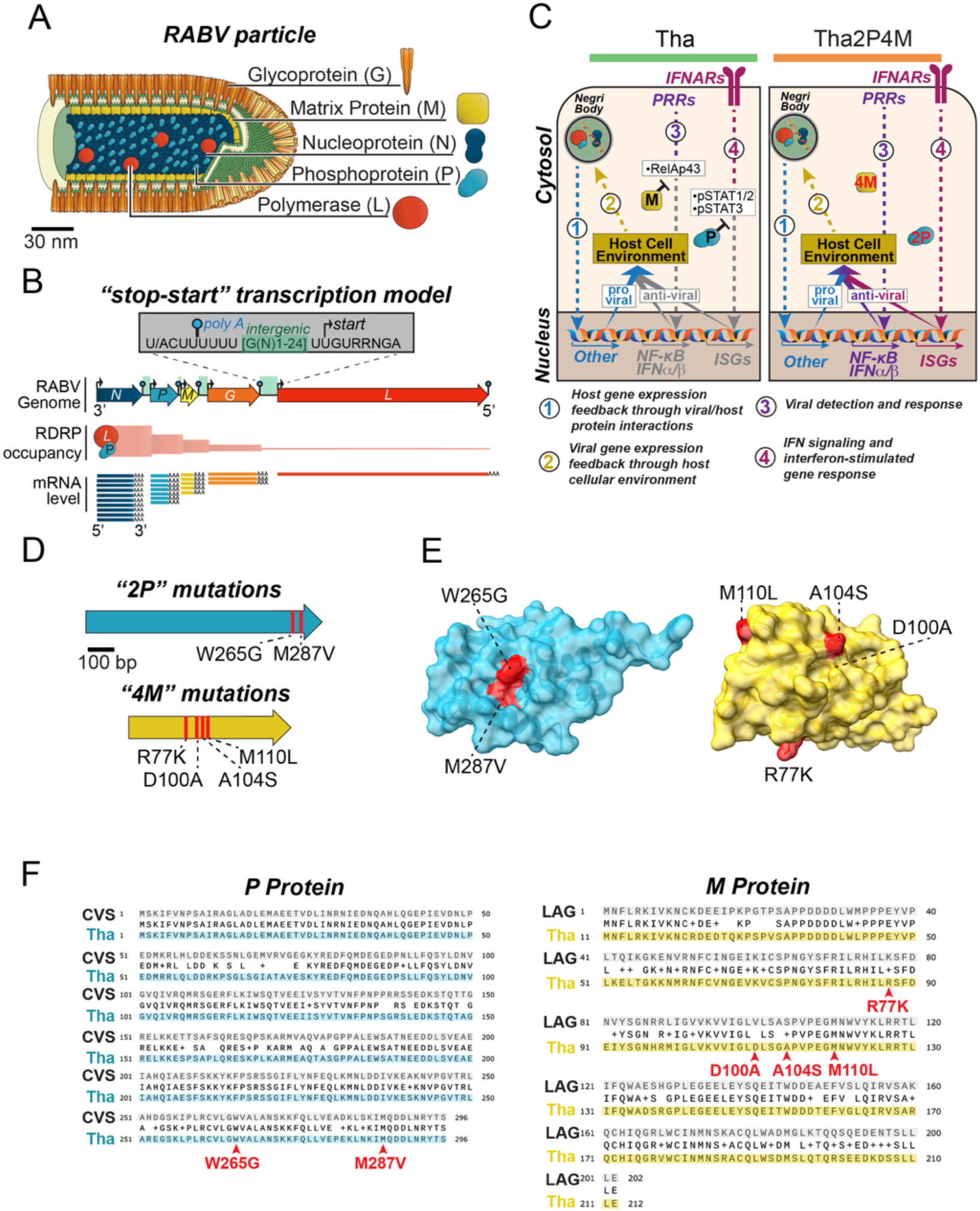
“2P4M” mutations in Tha RABV P and M proteins disinhibit innate immune detection and responses in host cells by neutering negative regulation of NF-kb and IFN-stimulated gene transcriptional factors. **(A)** Schematic illustrations of the RABV particle (modified from^2^). **(B)** Schematic of the RABV genome and accepted “start-stop” model of transcription (afteR^2^). RABV genes are expressed at levels that reflect genome order due to the occupancy of the RNA-dependent RNA Polymerase (RNDP) complex composed of the polymerase (L) and co-factor P. RNA sequence motifs responsible for the stop/polyadenylation (“poly A”) and transcriptional onset (“start”) functions of the RNDP are shown. **(C)** Schematic illustrating known mechanisms of RABV P and M inhibition of viral detection and response in Tha (*left*) or Tha2P4M (*right*) infection along with basic pathways (numbered below) through which viral/host interactions and inhibited/disinhibited innate immune signaling could affect viral and host cellular processes. Germline-encoded pattern recognition receptors (PRRs) are viral sensors (such as toll-like receptors (TLRs); retinoic acid-inducible gene I (RIG-I) like helicases (RLRs); and nucleotide-binding oligomerization domain-like receptors (NOD-like receptors) that induce NF-kB and IFN pathways upon activation. **(D)** Gene locations and amino acid alterations of six key point mutations that restrict innate immune inhibition by P (“2P”) and M (“4M”) proteins. **(E)** Protein structures of lyssavirus P (RABV CVS strain^106^) and M (Lagos bat virus^107^) highlight the outward-facing locations of mutant residues in P and M (red). P mutations cluster around the “W hole” (after^34^); M mutations affect three spatially-distinct interaction sites.**(F)** Amino acid-based protein alignments comparing Tha RABV P and M proteins versus those protein sequences from the solved structures shown in (**E**)(CVS P^106^ (PDB Accession 1vyi) and Lagos Bat M^107^ (PDB Accession 2w2s). The sites of the “2P” and “4M” mutations are shown in red.

**Supplemental Figure 2.**
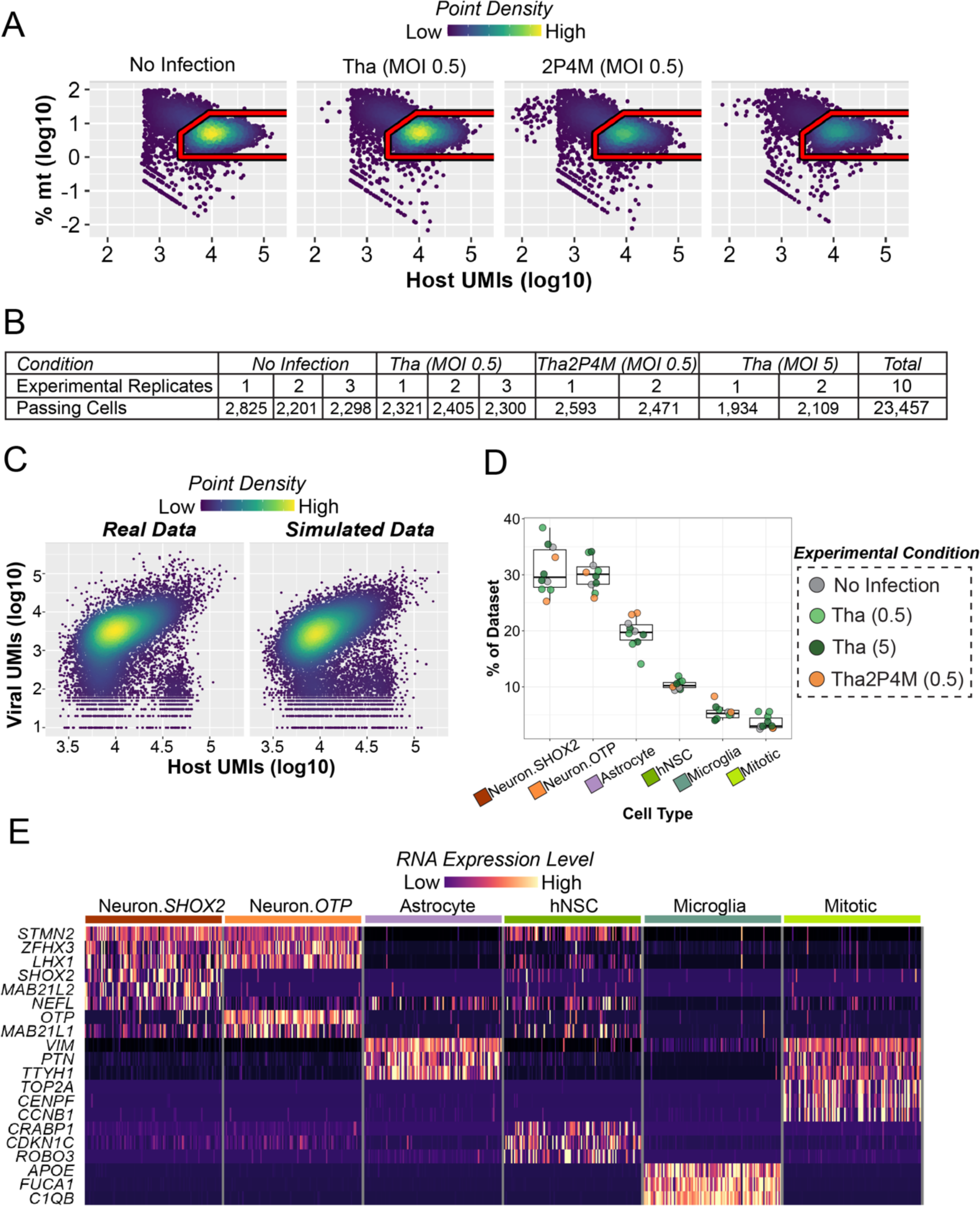
Quality-control metrics for scRNA-seq analysis and expectation-maximization approach for distinguishing RABV infected and uninfected bystander cells from human brain co-cultures. (A) Quality-control metrics for inclusion of scRNA profiles. Density scatter plots compare the size of RNA libraries (based on total human UMI counts) versus the fraction of RNA derived from the mitochondrial genome. The red polygons define high-quality RNA libraries included for downstream analysis. Colored heat map indicates point density. (B) Table summarizing the number of RNA profiles for each replicate across experimental conditions. (C) Density plot-based comparison of the real versus simulated data of viral versus total UMI counts, demonstrating the accuracy of the model used for Expectation-Maximization identification of RABV infected versus uninfected cells (**Methods**). (D) Box plot of cell type percentages across scRNA-seq datasets. Replicate-specific cell type percentages are shown with dots color-coded by experimental condition. (E) Heat map depiction of expression levels of select marker genes for transcriptional clusters associated with major human brain cell types. Colored heat map indicates expression level. Cells are downsampled by cell type for visualization.

**Supplemental Figure 3.**
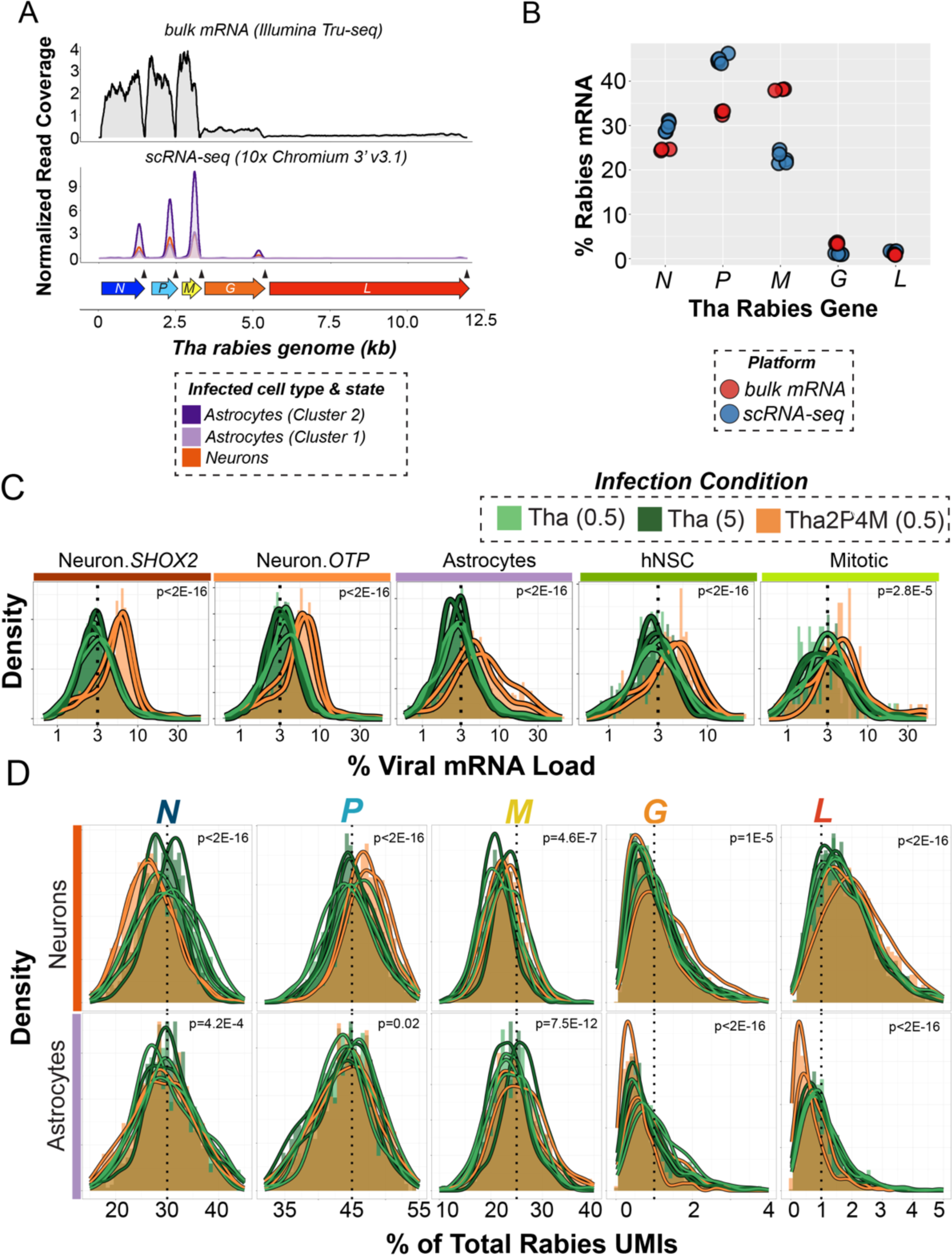
RNA expression analysis of Tha RABV genes. **(A)** Comparison of normalized sequencing read coverage across the Tha RABV genome for bulk RNA sequencing (*top*) following poly A capture and enrichment^35^ versus subsets of n=10 scRNA-profiles associated with Neurons and Cluster 1 and Cluster 2 Astrocytes (*bottom*)(Figure 4A; **Methods**). A schematic of the Tha RABV genome is shown at bottom and the locations of polyadenylation signals are indicated with arrow heads. **(B)** Comparison of relative expression levels for each Tha RABV gene based on bulk RNA sequencing (red; three biological replicates) versus the scRNA-seq analysis used in this study (blue; five total biological replicates, n=3 Tha(0.5) and n=2 Tha(5) aggregated across all cell types). Range mean % viral mRNA, bulk: *N* = 24.3-24.6, *P* = 32-33, *M* = 38-38.2, *G* = 3.3-3.7, *L* = 0.8-1.0, scRNA: *N* = 29-31, *P* = 44-46, *M* = 21-22, *G* = 0.9-1.2, *L* = 1.1-1.2. **(C)** Combined histogram and density plots of viral mRNA load stratified by brain cell types and color-coded by experimental replicate. P values reporting differences in distributions by experimental condition are reported in each panel (Kruskal-Wallis tests). **(D)** Density plots of relative RNA levels for each of the five Tha RABV genes color-coded by experimental replicate and stratified by both gene and neuron (*top*) or astrocyte (*bottom*) identity. P values reporting differences in gene expression distributions by experimental condition are reported for neurons and astrocytes in each panel (Kruskal-Wallis tests).

**Supplemental Figure 4.**
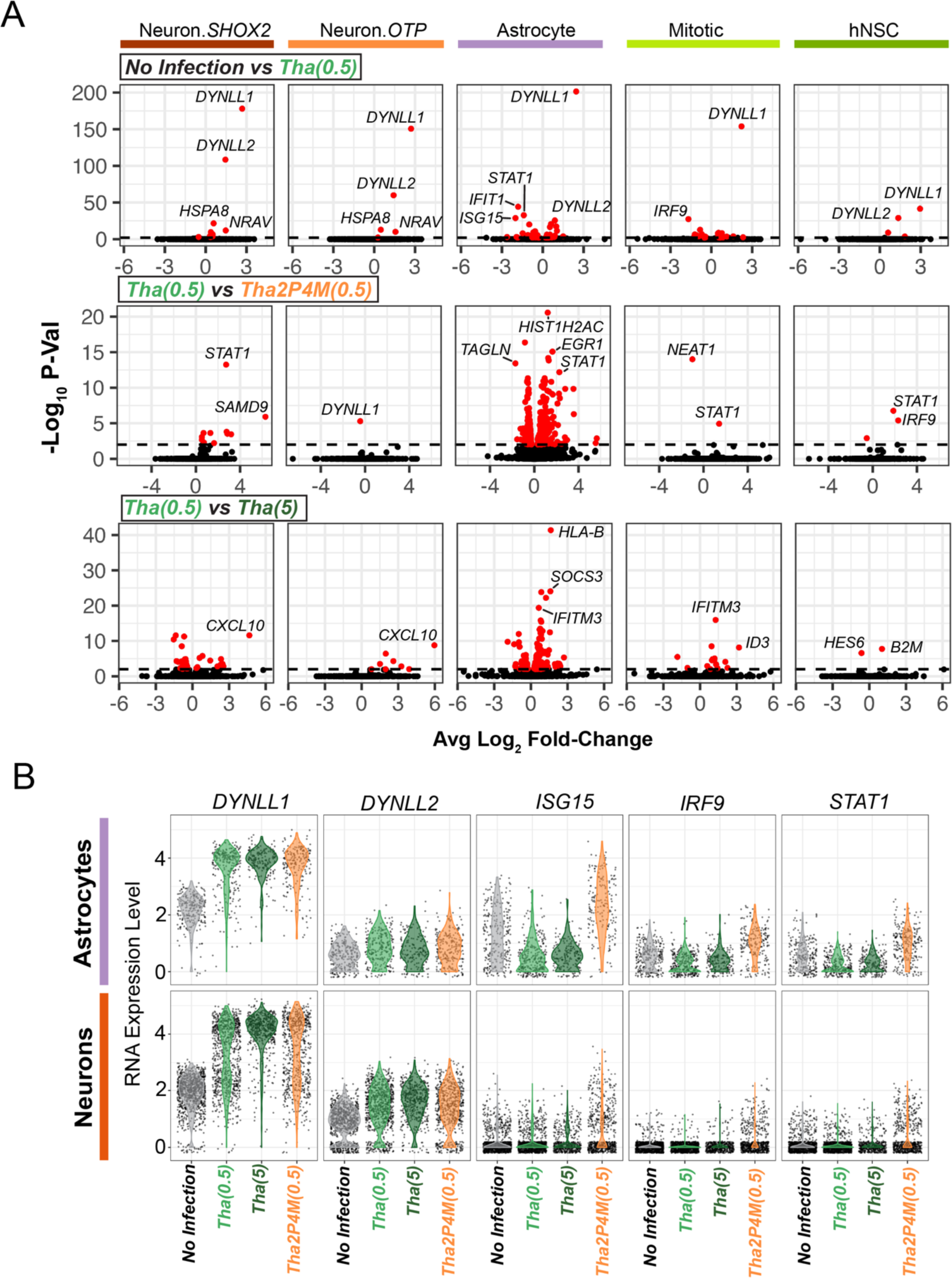
Host cell gene expression responses to different RABV infection conditions. **(A)** Volcano plots of genome-wide expression differences stratified by cell type and experimental condition (*top row*, No Infection versus Tha(0.5); *middle row*, Tha(0.5) versus Tha2P-4M(0.5); *bottom row*, Tha(0.5) versus Tha(5)). Select genes with significant differences (red, p < 0.01) are labelled for clarity. **(B)** Combined violin dot plots stratified by astrocytes (*top*) or neurons (*bottom*) illustrating RNA expression levels for key genes stratified by experimental condition. (RNA levels are shown as after log-normalization and centering following the default Seurat workflow; **Methods**).

**Supplemental Figure 5.**
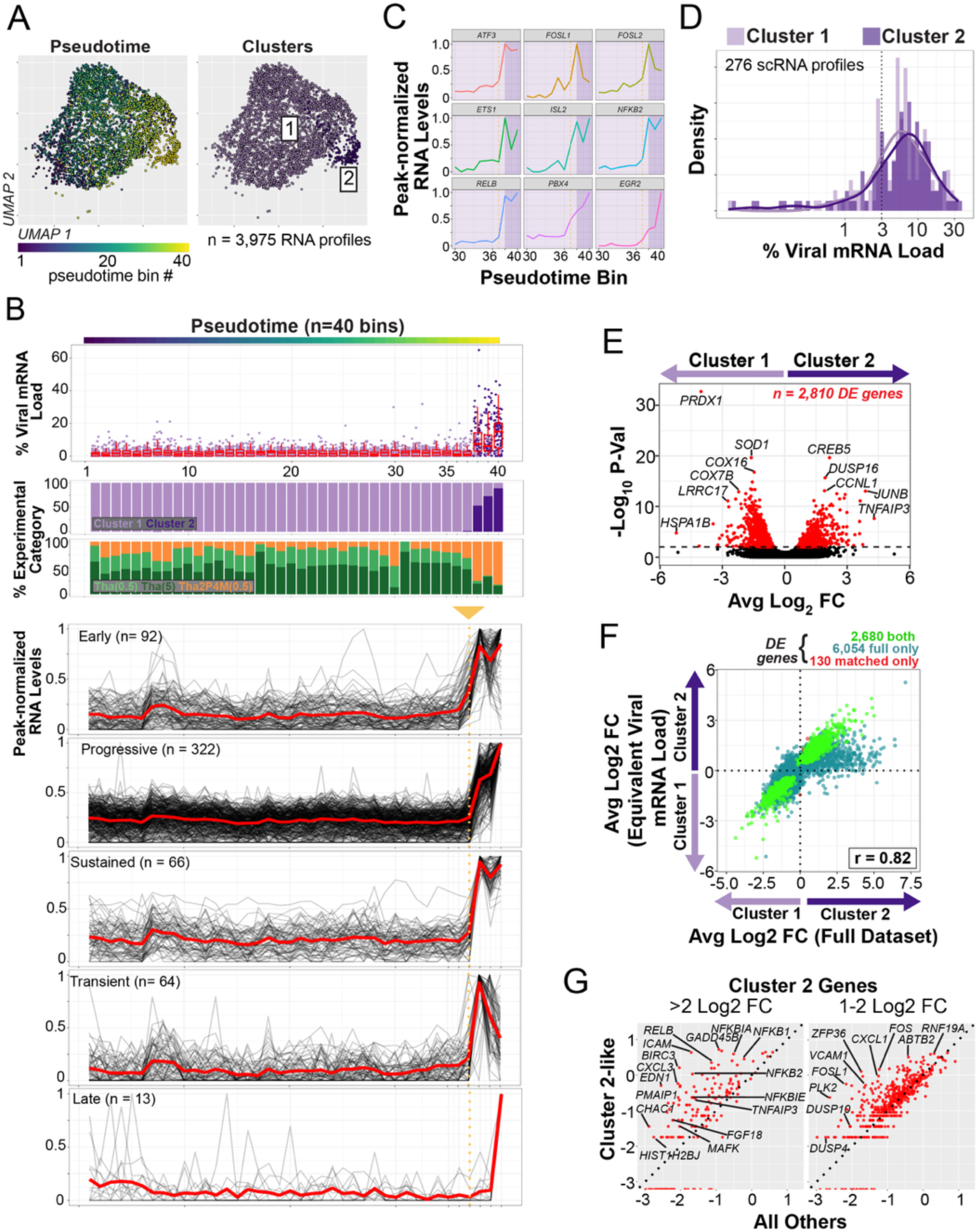
Large-scale gene expression changes driving pro-viral Cluster 2 astrocyte identities precede and are largely independent of viral mRNA load. **(A-C)** Pseudotime trajectory analysis of astrocyte scRNA profiles performed using Monocle3 **(Methods)**. **(A)** UMAP embeddings of astrocytes. *Left*, after binning into n=40 pseudotime groups and color-coded by a heat map. *Right,* Cluster 1 and Cluster 2 identities from the Seurat-based clustering analysis (Figure 4A). **(B)** Linear plots of astrocyte experimental or expression properties ordered by psuedotime bin. *Top*, viral mRNA load scores of each astrocyte (dots) color-coded by cluster identity along with bin-specific box plots (red). *Middle*, bar plots displaying the proportion of astrocytes by cluster or experimental condition. *Bottom*, line plots of peak-normalized gene-specific expression signatures across pseudotime bins stratified by temporal expression pattern (black). Mean expression values across all genes in each category are shown in red. Dotted gold line and arrow mark the bin (number 37) preceding the increase in viral mRNA load and pro-viral Cluster 2 identities. “Early” genes exhibit clear increases in expression that precede bin 37 in pseudotime (**Methods**). “Progressive”, “Sustained”, “Transient” and “Late” categories describe distinct expression signatures between bin 38 and 40 (**Methods**). **(C)** Peak-normalized expression signatures for n=12 transcription factors, selected as examples from the n=24 “early” genes identified as transcriptional regulators (belonging to the “chromatin” (GO:0000785) “cell component” Gene Ontology category (*ATF3, CENPA, CREB5, CSRNP1, EGR2, ETS1, FOSL1, FOSL2, FOXS1, HEY2, IRF1, ISL2, JADE1, JUNB, KLF5, MAFF, MEF2C, NFATC2, NFKB2, PBX4, RELB, TEAD4*). “Chromatin” genes were two-fold enriched in the “early” gene set (n=22 of 92; 24%) as compared to genes across all other categories in (**A**) (n=86 of 688; 12.5%)(p = 0.0008, Chi Square Test). Line plots for each gene are named and color-coded. Only pseudotime bins 30-40 are shown to highlight the Cluster 1 to Cluster 2 transition. The dark purple background highlights bins 38-40 in which pro-viral astrocytes emerge (**B**). **(D)** Combined histogram and density plot of viral mRNA load distributions for Cluster 1 and 2 after downsampling each cluster to n=276 scRNA profiles with equivalent viral mRNA load distributions (**Methods**). **(E)** Volcano plots of gene expression differences across viral mRNA load downsampled Cluster 1 and 2 with select genes labelled (red, p < 0.01). **(F)** Scatter plots comparing log2 fold expression level changes for genes significantly altered (p < 0.01) across Cluster 1 or Cluster 2 in the full dataset or a dataset in which viral mRNA load scores were downsampled to an equivalent distribution in Cluster 1 and 2 (**Methods**). Differentially expressed genes are color-coded by membership (shared, green; full dataset only, cyan; equivalent viral load only, red). **(G)** Scatterplots of normalized metacell expression values for the 196 genes used to identify “Cluster 2-like” profiles (Identification Gene Set: p < 0.01 and >2 log fold-change; *left*) and for an independent set of n=521 gene RNAs with enriched expression in Cluster 2 astrocytes (Validation Gene Set: p < 0.01 and 1-2 log fold-change; *right*)(**Methods**). Metacells are composed of n=38 “Cluster 2-like” nuclei RNA profiles versus all other astrocyte nuclei profiles in the NPH dataset (“All Others”; n=82,596). Total UMI counts for both NPH gene sets were significantly enriched in “Cluster 2-like” NPH astrocyte profiles at levels that matched expectation from Cluster 2 (Identification Gene Set, 2.8 fold enriched; Validation Gene Set, 1.2 fold enriched; p<2.2E-16, Chi Square Test).

**Supplemental Figure 6.**
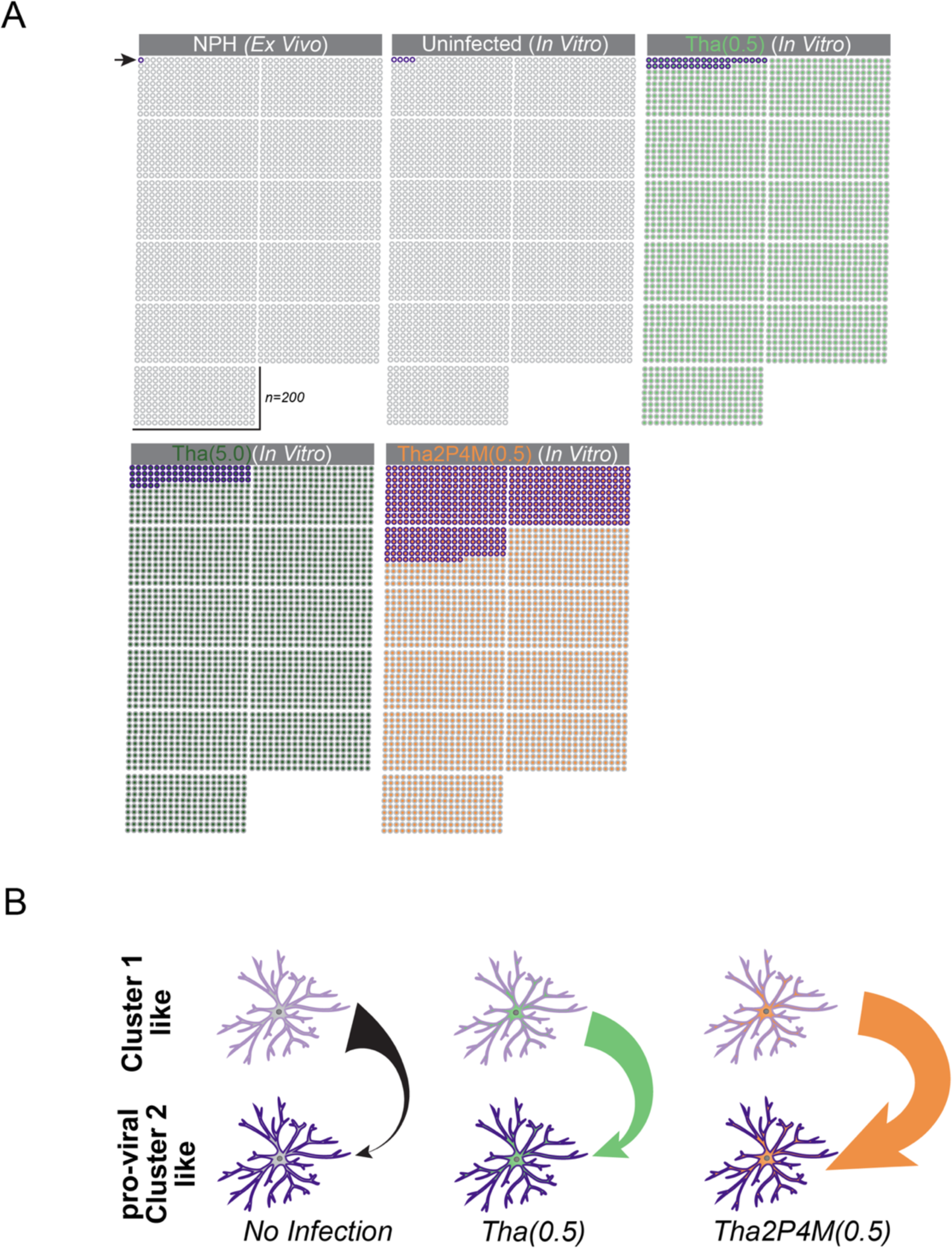
Schematic of abundances and working model of pro-viral astrocyte state transition. **(A)** Schematic approximating the abundance of astrocytes in the Cluster 2 (or Cluster 2-like) pro-viral state across experimental conditions estimated from the NPH and Tha datasets. Circles representing 2,200 astrocytes are shown. The experimental state is indicated by the dot fill, while the Cluster 2/Cluster 2-like state is indicated by a purple outline (beginning at top left, as indicated by the arrow). **(B)** Cartoon model illustrating the relative abundances and Cluster 1-like or pro-viral Cluster 2 like astrocytes in different RABV conditions. Arrowhead and tail width indicate the relative proportion of each astrocyte state, with larger widths indicating higher proportions. Without infection, pro-viral Cluster 2 like astrocytes are present, but rare. RABV infection increases the likelihood of Cluster 1-like astrocytes transitioning to a pro-viral like state. When the 2P4M mutations are present, the likelihood of this transition is much higher. It is unknown whether pro-viral astrocytes can transition back to a more Cluster 1 like state.

**Supplemental Figure 7.**
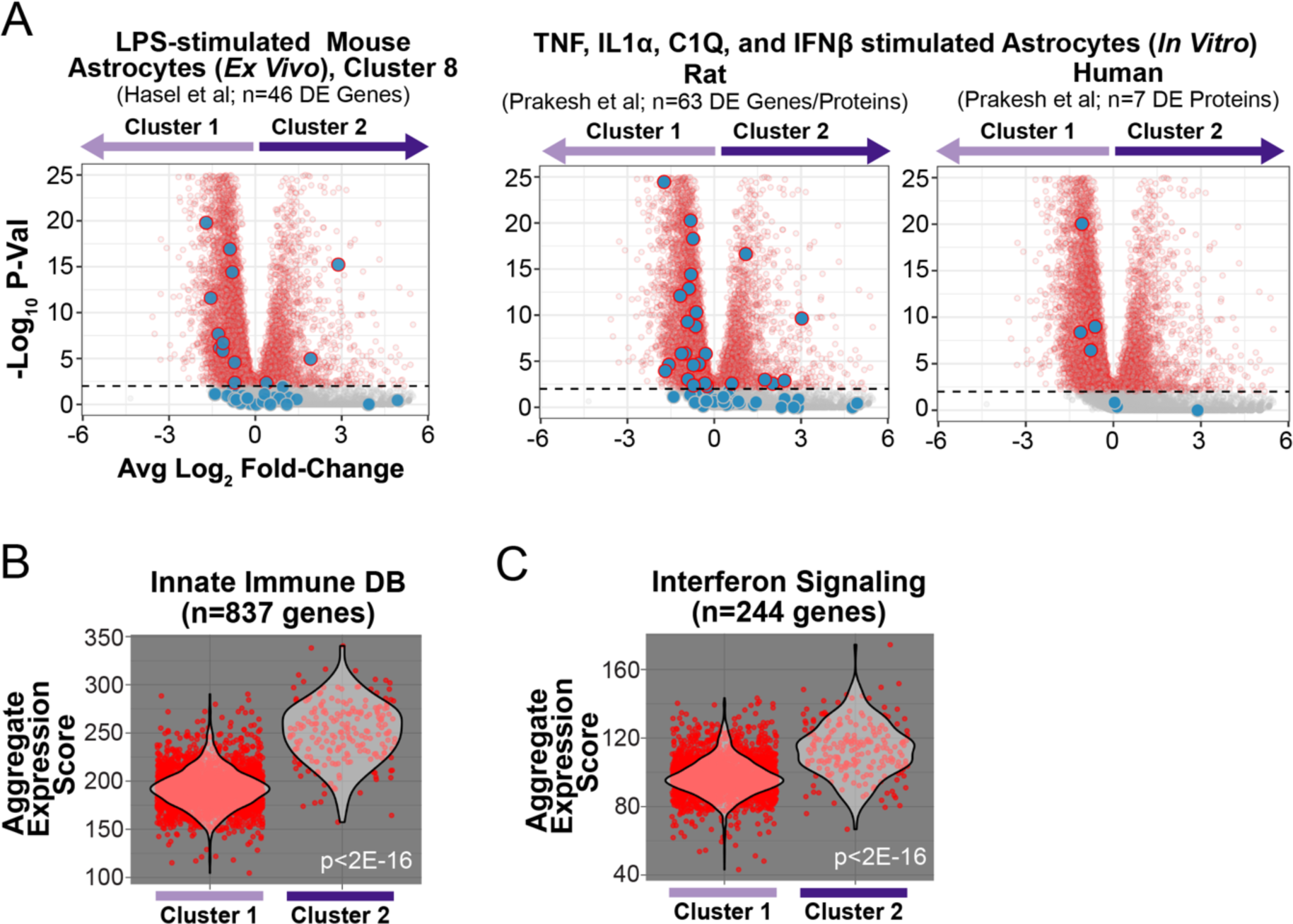
Gene expression signatures observed in pro-viral Cluster 2 astrocytes are not observed in previous descriptions of “reactive” rodent or human astrocytes. **(A)** Plots of differentially expressed RNAs or proteins from previous studies plotted on volcano plot of depicting expression differences across Cluster 1 and Cluster 2 astrocytes. The size and color of dots represents the intersection of differentially expressed RNAs or proteins from previous studies with those RNAs differentially expressed across Cluster 1 or Cluster 2. Large blue dots correspond to previously described as differentially expressed gene/protein sets (p < 0.01); those with red outlines are also significant in the Cluster 1 versus Cluster 2 comparison (p < 0.01). Smaller dots correspond to those genes observed as differentially expressed across Cluster 1 and Cluster 2 (red, p < 0.01; grey, p >= 0.01) but are not observed as differentially expressed in previous studies. *Left*, 46 genes differentially expressed following scRNA-seq analysis of mouse cortical astrocytes *ex vivo* with an IFN activated gene signature following Lipopolysaccharides (LPS) stimulation^81^. Primary rat (*middle*) or iPSC-derived human (*right*) astrocytes cultured *in vitro* and stimulated with TNF, IL1α, C1Q and IFNβ to induce an interferon-responsive state^80^. The 63 rat genes show concordant changes from bulk RNA-seq and proteomic analysis; the 7 human genes identify alterations based on proteomic analysis of secreted proteins (bulk RNA-seq was not performed). (**B,C**) Aggregate expression scores across Cluster 1 and pro-viral Cluster 2 astrocytes for those n=837 genes annotated in the InnateImmuneDB database^82^ (**B**) and n=244 human genes annotated in the “Interferon Signalling” REACTOME database category (R-HSA-913531)(**C**)(Wilcoxon Test).

**Supplemental Figure 8.**
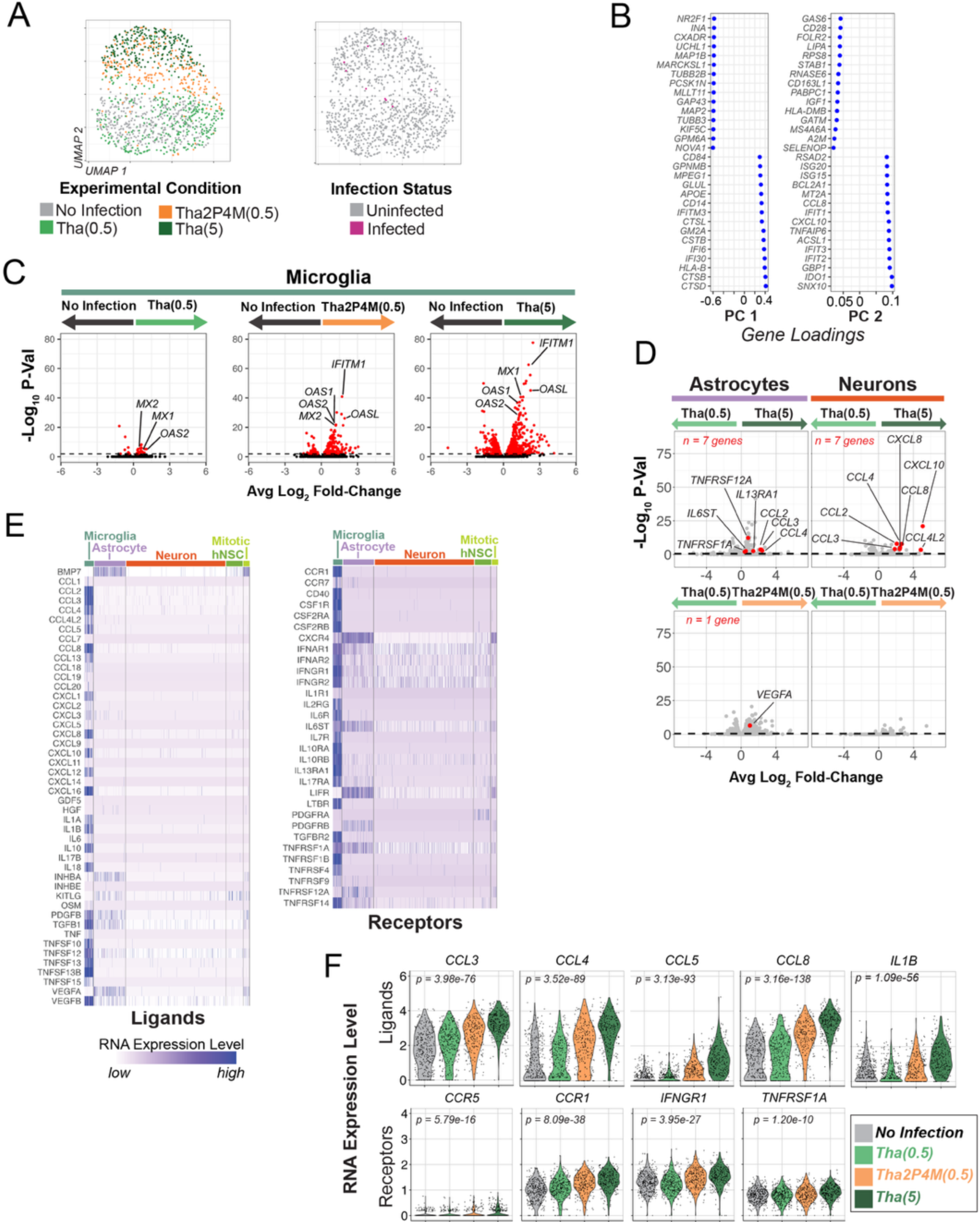
Microglia transcriptional response to Tha RABV infection of other cell types. (A) UMAP embedding plot of microglia scRNA profiles color-coded by experimental condition (*left*) and infection status (*right*). (B) Gene principal component loading scores for the top and bottom n=15 genes for PC 1 and PC 2. PC 1 genes suggest contamination of neuronal transcripts, while PC 2 genes represent genes transcriptionally regulated by interferon signalling. (C) Volcano plots of gene expression differences with select genes labelled (red, p < 0.01). The corresponding fold-change values for these analyses are shown in Figure 6C. (D) Volcano plots displaying gene expression differences across experimental conditions for astrocytes and neurons (*top*, Tha(0.5) vs. Tha(5)(*top*); *bottom*, Tha(0.5) vs. Tha2P4M(0.5)). KEGG cytokine-cytokine receptor interaction pathway (M9809) genes with (p < 0.01) are shown in red. All other genes in grey. (E) Gene expression heatmap of all cytokine ligand and receptor genes (KEGG cytokine-cytokine receptor interaction pathway M9809) (ligands, *left*; receptors, *right*) across cell types including all experimental conditions. (F) Violin plots of microglia expression across experimental conditions for select cytokine ligands (*top*) and receptors (*bottom*) with known roles in NF-κB signalling (effect of experimental condition assessed with a one-way ANOVA, α < 0.01). Example ligands and receptors show graded expression levels that are similar to microglia ISGs across experimental conditions (Figure 6E).

## Supplemental Table

**Supplemental Table S1** contains all differential gene expression testing results reported in the manuscript.

## STAR Methods

### KEY RESOURCES TABLE

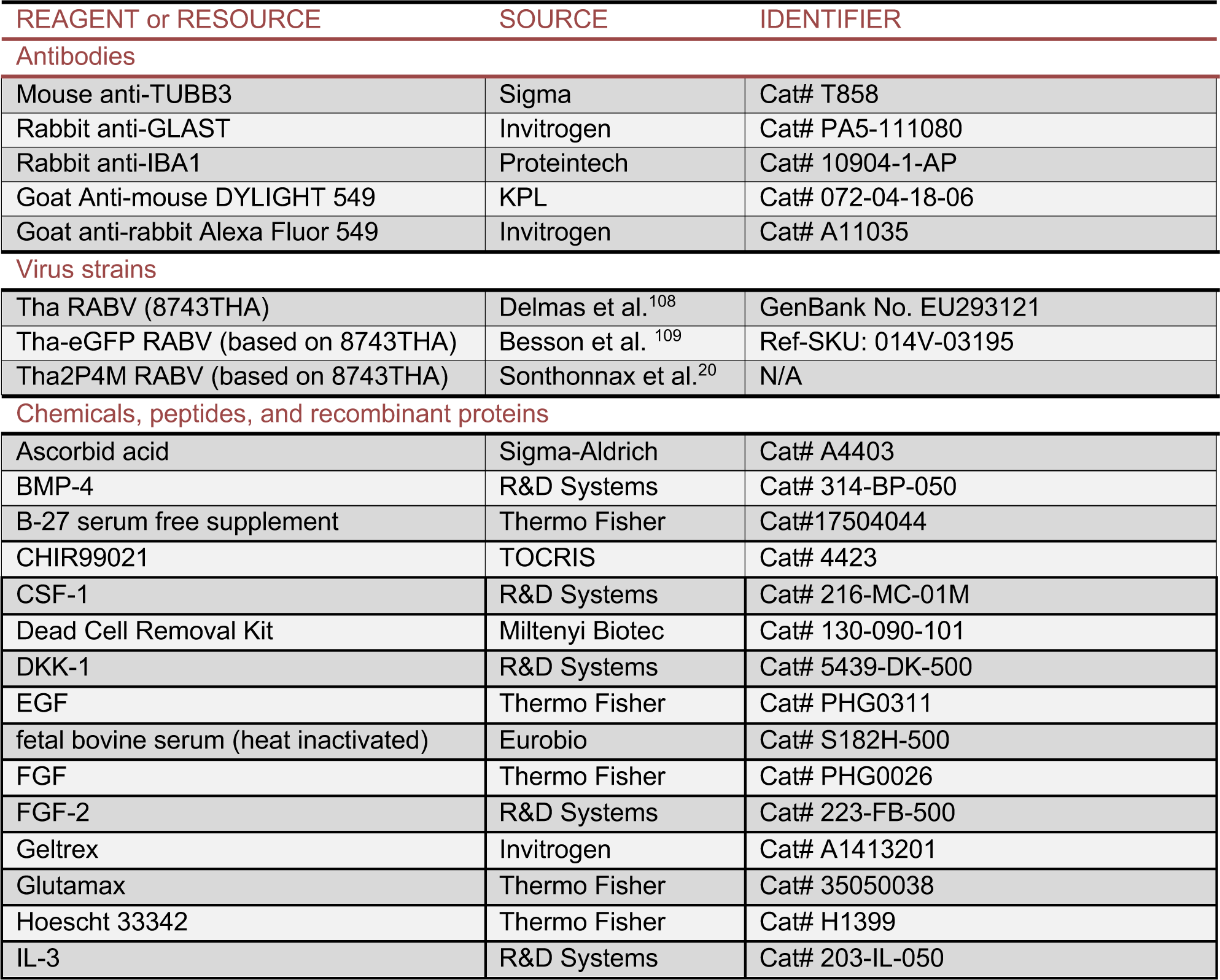
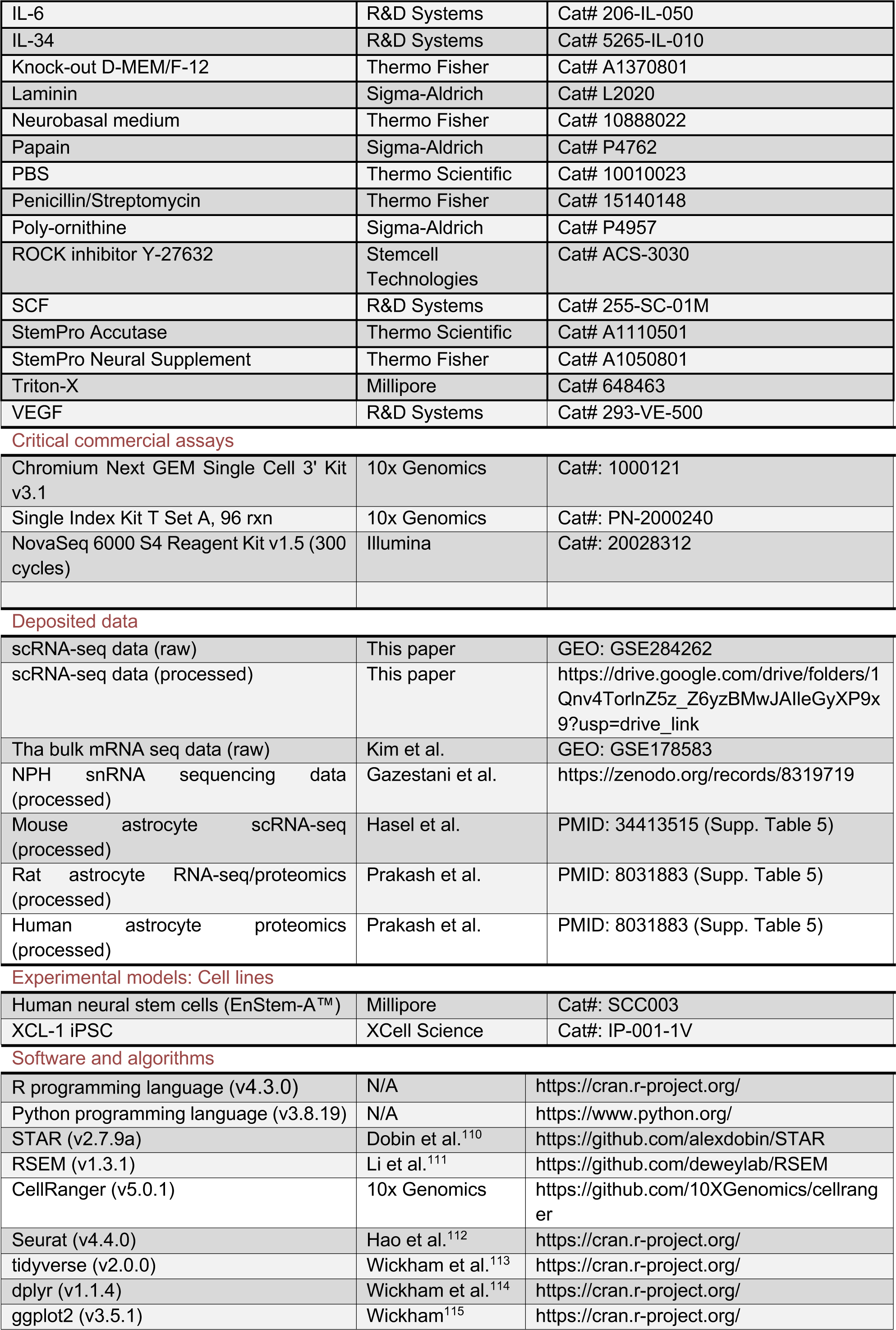
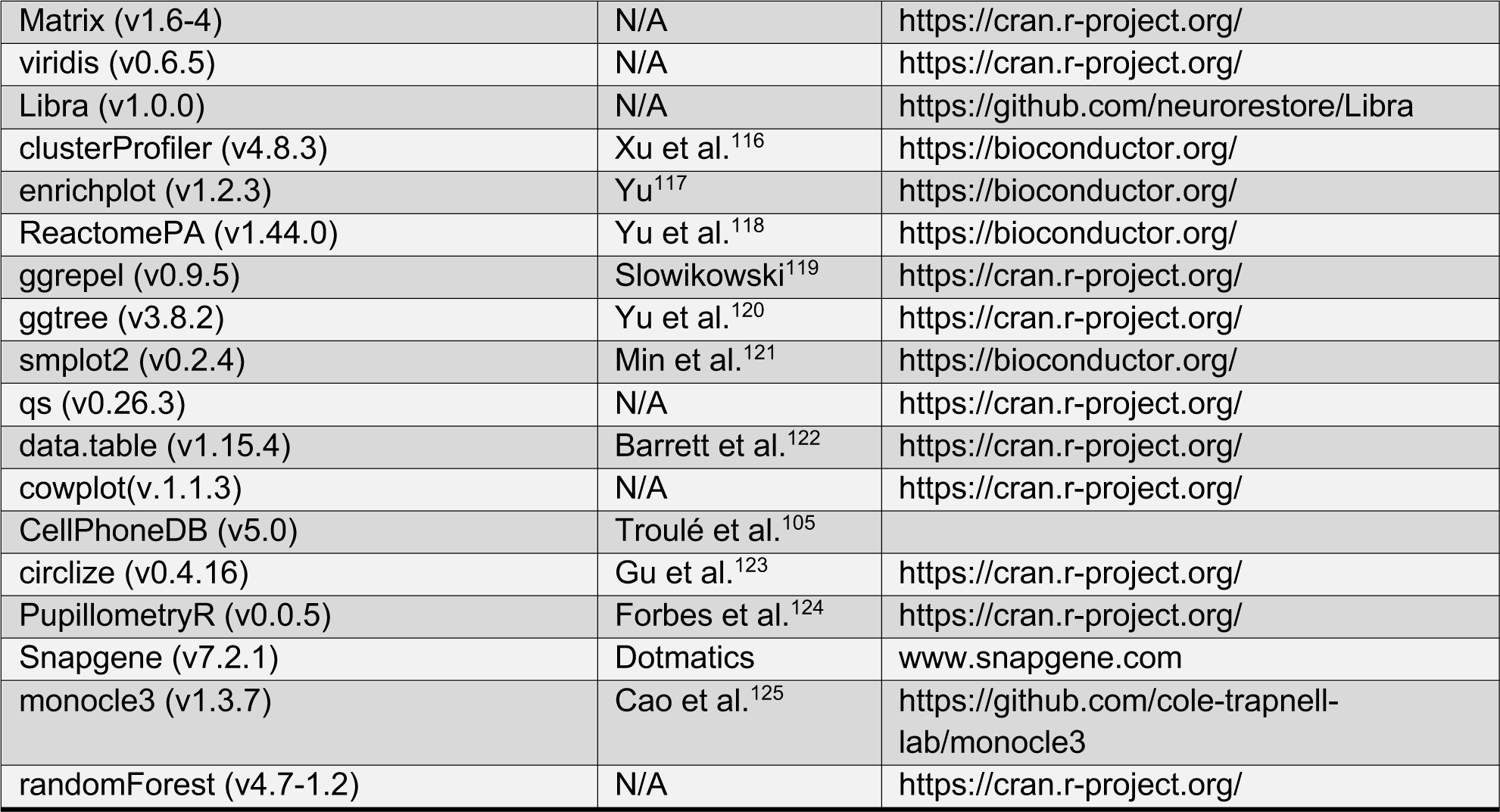

### RESOURCE AVAILABILITY

#### Lead contact

Further information and viral reagent requests will be fulfilled by lead contacts Dr. Arpiar Saunders and Dr. Hervé Bourhy.

#### Materials availability

Tha RABV reagents are available from Dr. Hervé Bourhy with a completed MTA and appropriate lab safety and import restrictions.

#### Data and code availability

The scRNA-seq datasets generate for this study are available from Gene Expression Omnibus Accession GSE284262. All analysis scripts and processed data are available (https://drive.google.com/drive/folders/1Qnv4TorlnZ5z_Z6yzBMwJAIleGyXP9x9?usp=drive_link).

### EXPERIMENTAL MODEL AND SUBJECT DETAILS

#### Ethics Statement

All Tha RABV *in vitro* infections were performed under BSL3 containment in accordance with the biosafety protocols developed and approved by the Pasteur Institute.

#### Human brain cell type co-culture

##### Human neural stem cell culture and differentiation

Human neuronal stem cells (hNSC, EnStem-A™ SCC003, Millipore) were amplified on plates previously cultured with geltrex (A1413201, Invitrogen) in complete neural stem cell (CNSC) medium containing Knock-out D-MEM/F-12 (A1370801, Thermo Fisher), 2 mM Glutamax (35050038, Thermo Fisher), 2% StemPro Neural Supplement (A1050801, ThermoFisher), 20 ng/ml human FGF (PHG0026, Thermo Fisher), and 20 ng/mL human EGF (PHG0311, Thermo Fisher) at 37°C and 5% CO_2_. For differentiation, hNSC cells were seeded at a density of 1.5×10^5^ cells/cm^2^ on plates previously coated with poly-ornithine (P4957, Sigma-Aldrich) and laminin (L2020, Sigma-Aldrich). One day after seeding, the medium was changed to differentiation medium containing neurobasal medium (10888022, Thermo Fisher), 2 mM Glutamax (35050038, Thermo Fisher), 2% B-27 serum free supplement (17504044, Thermo Fisher), 200 *μ*M Ascorbid acid (A4403, Sigma-Aldrich) and 1x Penicillin/Streptomycin (15140148, Thermo Fisher). Human NSC were differentiated for one week before hiMacs were added to the culture. Half of the medium was changed every third day.

#### Differentiation of hiPSCs to hiMacs

Human induced macrophage-like cells (hiMacs) were generated from iPSCs according to a published method described by Takata *et al*^126^. In short, 5 ng/mL BMP-4 (314-BP-050, R&D Systems), 50 ng/mL VEGF (293-VE-500, R&D Systems), 2 *μ*M CHIR99021 (4423, TOCRIS) induced mesoderm specification of human iPSC colonies (XCL-1) and hemangioblast-like cell formation (36). Replacement of CHIR99021 by 20 ng/mL FGF-2 (223-FB-500, R&D Systems), followed by maintenance in 15 ng/mL VEGF and 5 ng/mL FGF-2 induced differentiation into the hematopoietic lineage according to a modified protocol from Grigoriadis *et al.* (37). Wnt signalling was inhibited and hematopoietic stem cells were matured by incubation with 50 ng/ml SCF (255-SC-01M), 10 ng/mL FGF-2, 20 ng/mL IL-3 (203-IL-050, R&D Systems), 10 ng/mL IL-6 (206-IL-050, R&D Systems), 10 ng/mL VEGF and 30 ng/mL DKK-1 (5439-DK-500) for 6 days and with 50 ng/mL SCF, 10 ng/mL FGF-2, 20 ng/mL IL-3 and 10 ng/mL IL-6 for 4 days. To promote terminal differentiation of hiMacs, cells were cultured with 50 ng/mL CSF-1 (216-MC-01M, R&D Systems) for 14 days (38–40).

#### Differentiation of hiMacs to hiMicros

After hiMacs generation, hiMacs were co-cultured with 70 – 80 % confluent one week old (day 0 = induction of neural induction) hiNeurons and hiAstrocytes for three consecutive weeks. In detail, 7 days after the induction of neural differentiation, 1×10^4^ cells/cm^2^ hiMacs were added to the 1.5×10^5^ cells/cm^2^ hiNeurons and hiAstrocytes. Half of the culture medium was changed every third day supplemented with 50 ng/mL recombinant human CSF-1 (216-MC-010, R&D Systems) and 50 ng/mL human IL-34 (5265-IL-010, R&D Systems). Cells were co-cultured for three consecutive weeks before infection experiments were started.

#### Tha RABVs

Thailand virus, referred to as 8764THA (Genbank No. EU293111) is a field strain of RABV isolated from the brain of a Thai patient who died of RABV after being bitten by a rabid dog (EVAg collection, Ref-SKU: 014V-02106) (29). This virus was further adapted to cell culture on BSR cells (a BHK-21 clone, kindly provided by Monique Lafon, Institute Pasteur, Paris) (32), consequently called 8743THA (Genbank No. EU293121) (29) and in this manuscript referred to as “Tha”. The recombinant Tha2P4M RABV harbors the same genetic background as Tha apart from bearing two point mutations in the viral P-protein (W265G, M287V) and four mutations in the viral M-protein (R77K, D100A, A104S, M110L) which were previously described (15,21,33). To validate viral infection, recombinant viral Tha-eGFP and Th2P4M-eGFP constructs were used which were generated by cloning sequences of eGFP (recovered from pEGFP-C1 plasmid, Promega) into the genetic sequence of Tha (34) and Th2P4M, respectively. Viral strains were sequenced before used for infection experiments. Viral titers were determined by virus titration on BSR cells via staining of the viral nucleoprotein (5100, Sigma-Aldrich).

### METHOD DETAILS

#### Visualizing 2P4M mutations on P and M protein structures

ChimeraX (v1.8)^127^ was used to highlight the P protein (W265G, M287V) substitutions on the structure of the P protein from RABV CVS-11 strain (Protein Data Bank Accession: 1vyi) and four mutations in the M protein (R77K, D100A, A104S, M110L) on the M protein from Lagos bat virus (Protein Data Bank Accession: 2W2S). Amino acid alignments (using the CLUSTALW algorithm implemented in SnapGene (v7.2.1)) confirmed the conserved site positions of each mutation across lyssavirus species and strains (**Figure S1F**).

#### Tha RABV infection conditions

After four weeks of differentiation *in vitro*, co-cultured human brain cells were subject to Tha RABV infection through bath-application of Tha RABV (diluted in 500 ul of differentiation media) versus mock infections (500 ul of differentiation media alone) for two hours at 37°C and 5% CO_2_. To visualize infected cells and confirm identities of infected cell types, Tha RABV expressing GFP (Tha-eGFP, MOI 0.5) was used following post-hoc immunofluorescence analysis. To generate co-cultures upstream of single-cell RNA sequencing, infections were performed with Tha (MOI 0.5 or 5.0), Tha2P-4M (MOI 0.5) or “No Infection” (mock infection) were performed.

#### Immunofluorescence-based identification of Tha RABV infected cell types

Forty-eight hours post-infection, co-cultures were fixed using 4% PFA for 15 minutes at room temperature. After fixation, cells were then washed with PBS (10010023, Thermo Scientific) and permeabilized using 0.5% triton X-100 (648463, Millipore) for 10 minutes. Immunolabeling was performed with primary and secondary antibodies (Key resources table) using the following staining conditions: Cells were stained with the primary antibody diluted in PBS (10010023, Thermo Scientific) supplemented with 5% heat-inactivated fetal bovine serum (S182H-500, Eurobio) overnight at 4°C according to the manufacturer’s instructions. Subsequently, cells were washed using PBS (10010023, Thermo Scientific), followed by staining with the secondary antibody for 30 minutes at 37°C according to the manufacturer’s instructions. Immunolabeled cultures were washed using PBS (10010023, Thermo Scientific) and imaged using the Opera Phenix^TM^ High Content Screening System (Perkin Elmer) using standard conditions.

#### Single-cell RNA sequencing data generation and processing

Forty-eight hours after infection, cells were dissociated using StemPro Accutase (A1110501, Thermo Scientific) supplemented with 50 U/mL papain (P4762, Sigma Aldrich). In detail, cells were incubated in 400 *μ*l dissociation solution for 10 minutes at 37°C. After 10 minutes, cells were resuspended in neurobasal medium (10888022, Thermo Fisher) supplemented with 2% heat-inactivated fetal bovine serum (S182H-500, Eurobio) and 10 *μ*M ROCK inhibitor Y-27632 (ACS-3030, Stemcell Technologies). Cells were centrifuged at 1000 *rpm* for 5 minutes at 4°C. Viability was increased by removing dead cells via the Dead Cell Removal Kit (130-090-101, Miltenyi Biotec). In short, cells were resuspended in 100 *μ*l of dead cell removal beads. After 15 minutes of incubation, 400 *μ*l of binding buffer was added to each sample. Samples were added to the freshly washed LS columns (130-042-401, Miltenyi Biotec). Columns were washed with 4.5 mL binding buffer (Miltenyi Biotec) and the flow-through was collected. After the collection of viable cells, cells were centrifuged at 1000 *rpm* for 5 minutes at 4°C. Meanwhile, viable cells were counted and diluted to a final concentration of 1000 cells/*μ*l. 10,000 cells were loaded per condition using the Chromium Next GEM Single Cell 3ʹ Reagent Kit v3.1 (PN-1000128, 10x Genomics). The 10x libraries were sequenced at a depth of 40,000 reads per cell using the Illumina NovaSeq Instrument (Novogene). CellRanger (10x Genomics; v5.0.1) was used to align reads against a composite human (GRCh38) and Tha RABV genome. Digital Gene Expression (DGE) matrices were generated using reads mapped to gene exons.

#### scRNA-seq workflow for analysis of Tha RABV and human gene expression

Single-cell RNA-seq data was analyzed in two distinct ways to study human and viral gene expression in an independent manner and in a synergistic manner in which viral and human RNAs were associated with the same RNA profiles using experiment-specific cell barcodes. First, to study human gene expression patterns without the influence of Tha RABV RNAs, Tha RABV genes were removed to create human-specific DGEs. Human-specific DGEs across all experimental conditions were filtered based on quality control metrics (described below) and then processed using a standard workflow in Seurat (v4.4.0)^112^ to clustering RNA profiles, revealing cell type identities. Specifically, scRNA libraries were filtered to include those libraries with >2,500 UMIs and % of mitochondrial RNAs meeting custom thresholds based on library size (**Figure S2A**; range: 1-19.9 %). UMI counts in each library were then log-normalized (*NormalizeData(normalization.method = “LogNormalize”, scale.factor = 10000)*) and expression values for each gene were centered (*ScaleData()*). Clustering was then performed after variable gene selection (*FindVariableFeatures()* and *RunPCA()*), using n=15 principal components as input for shared nearest-neighbor graph construction (*FindNeighbors(dims=1:15)*). Clustering was then performed with the smart local moving algorithm (*FindClusters(resolution=0.12)*) and a UMAP embedding was generated (*RunUMAP(dims=1:15, min.dist=0.4))* for visualization. Cell type identities for each cluster were assigned by evaluating the expression patterns of well-established cell-type-specific marker genes for neurons, astrocytes, microglia, neural stem cells and mitotic cell (**Figure S2E**). Second, an “infection summary” file was generated to study Tha RABV gene expression at single-cell resolution. The summary infection file includes UMI counts for each Tha RABV gene as well as total Tha RABV and grand total UMIs (human + Tha). Cell type and infection-status identities (see below) were then assigned to the Tha infection file using experiment-specific cell barcodes for downstream analysis. Importantly, to avoid confounds of low-quality libraries, only those RNA profiles passing the quality-control metrics and included in the final Seurat analysis were included for analyses of Tha gene expression. After global clustering analysis, astrocyte and microglia cellular RNA profiles were independently analyzed in Seurat using a similar workflow. In each case, subclustering revealed minor groups of RNA profiles that exhibited gene expression signatures of other cell types or high-levels of nuclear lncRNAs and thus were removed as candidate “doublets” or low-quality cellular RNA profiles.

#### Identification of infected and uninfected cells using expectation-maximization

The observed Tha RABV viral RNA UMI counts were modelled as a mixture of two compound log-normal-binomial distributions, one for each of the populations of infected and uninfected cells. These compound distributions were also regressed by the total UMIs for each profile (i.e. library size), allowing the model to account for differences infectivity across cell type populations with varying library sizes. Specifically, letting *LS = library size (total Tha + human UMIs)*, *VL = viral load (% of UMIs from Tha)*, and *VC = viral counts (total Tha UMIs)*, for a given cell with index *i* we draw:

*VL_i ∼ logNormal (mu_offset + mu_coef * log (LS_i), sigma_offset + sigma_coef * log(LS_i))*

and then:

*VC_i ∼ Binom (VL_i, LS_i)*

where the parameters *mu_offset*, *mu_coef*, *sigma_offset*, and *sigma_coef* are learned separately for the populations of infected and uninfected cells. The conjugate prior distribution for the binomial distribution is the beta distribution, but we found the beta distribution to fit the expression data poorly, which we hypothesize is due to numerical instability arising from the extreme skew towards low viral load in our data. We chose instead to use a log-normal distribution as the prior distribution for VL, with the additional restriction 0 <= VL <= 1. (In practice, the missing/erroneous probability density from VL > 1 is negligible). Parameter learning was accomplished with a standard expectation-maximization algorithm, first randomly assigning each cell to one of two groups, which we will later categorize as infected or uninfected. For the maximization step, we fit a log-normal-binomial compound distribution (via maximum likelihood estimation) for each population of cells, as well as estimating the priors for each cell belonging to either group. To find maximum likelihood estimates for the log-normal-binomial distributions, we computed and maximized likelihood using the R language optimizer “optim” function, with “method = ‘Nelder-Mead’” and starting parameters ‘mu_off = −4, mu_coef = −0.1, sigma_off = 1.5, sigma_coef = 0’. The probability density for the log-normal-binomial distribution was computed by grid approximation (using 200 points evenly spaced between [-15, 0] and exponentiated). For the expectation step, each cell was assigned to the fitted model from the previous step that maximized its posterior probability. Then new compound log-normal-binomial distributions were fit to the reassigned groups of cells, and the cells recategorized, repeating this process until convergence (n=20 iterations).

#### Gene expression plotting

Gene expression levels were plotted using normalized, gene-centered data using functions from Seurat or custom plotting functions using the “ggplot2” (v3.5.1)^115^ R package.

#### Bulk and scRNA-seq analysis of Tha read alignment and gene expression

We re-analyzed bulk mRNA expression data from Tha RABV infection of BSR cells (Gene Expression Omnibus accession GSE178583) in addition to Tha RABV scRNA-seq data from human brain cell type co-cultures to compare Tha RABV gene expression levels and read alignment locations across sequencing modalities (**Figure S3A,B**). Bulk mRNA sequencing data was generated with the TruSeq Stranded mRNA library kit (Illumina). In the previous publication^35^, bulk mRNA data were aligned and analyzed with respect to only the human genome (reference GRCh38). Thus STAR (v2.7.9a)^110^ was used to realign the bulk mRNA fastq files to both the human and Tha genomes followed by RSEM (v1.3.1)^111^ to quantify Tha RABV gene expression levels using transcripts per million reads (TPM) values. Sequencing read coverage against the Tha genome was assessed for the bulk mRNA data and ten randomly selected cell barcodes from the scRNA-seq associated with the neurons, cluster 1 and pro-viral cluster 2 astrocytes. To do this, BAM files associated with these specific read sets were converted to bigwig files then read alignment coordinates were exported to csv files for analysis in R.

#### Tha RABV gene expression analysis from scRNA-seq data

Viral mRNA load scores were calculated for each cell’s RNA profile as the percentage of total UMIs (human + Tha) attributed to the five genes in Tha RABV genome (*N, P, M, G* and *L*). Relative expression of each Tha RABV gene was calculated based on the UMIs associated with each gene versus the total Tha RABV UMIs for each cellular RNA profile. To avoid biases driven by insensitive mRNA detection, analysis was limited to cellular RNA profiles with > 0 UMIs for *G* and *L* (the Tha genes with the lowest expression levels; **Figure S3B**). A one-way ANOVA was used to test for differences in Tha gene expression levels, while a two-way ANOVA was used to evaluate if cell type and experimental condition affected relative expression of each Tha gene. Pairwise testing of differences Tha gene expression by cell type and experimental condition was performed using the Kolmogorov–Smirnov test. P values are reported as uncorrected.

#### Differential expression testing

Human gene differential expression testing across experimental conditions or cell types was implemented in the “Libra” (v1.0.0) R package using the DESeq2 likelihood ratio test method (*Libra::run_de(de_method=“DESeq2”, de_type=“LRT”))*. All reported and analyzed p values are corrected for multiple hypothesis testing using the Benjamini-Hochberg method.

#### Gene Set Enrichment REACTOME analysis

Gene Set Enrichment Analysis (GSEA) and REACTOME Analysis were performed using the “clusterProfiler” (v4.8.3)^116^ and the ReactomePA(v1.44)^118^ R packages, respectively. GSEA assessed the “Biological Pathways” ontology using a gene-specific score based on multiplying the −log10(p value) * log2FC for each gene across Tha2P4M(0.5) versus Tha(0.5) astrocytes (*clusterProfiler::gseGO(ont = ‘BP’,OrgDb = ‘org.Hs.eg.db’, eps=1e-300)*). The ten Biological Pathways with strongest enrichment scores visualized using the enrichplot (v1.20.3) R package (*enrichplot::gseaplot2(= 1:10, ES_geom = “line”*). REACTOME analysis was performed on differentially expressed gene sets (p < 0.01) enriched in either Cluster 1 (n=2,137 genes) or Cluster 2 (n=5,887 genes) Astrocytes (p <0.01) (ReactomePA::enrichPathway(pvalueCutoff = 0.01). To discern structure within complex REACTOME results, hierarchical clustering was performed for the most enriched REACTOME categories Cluster 1 (p < 0.000001; n = 32 categories) and Cluster 2 (p < 0.01; n = 84 categories) based on similarities of gene driving each categorical enrichment. These results were visualized using the treeplot() function from the enrichplot (v1.20.3) R package. REACTOME p values are computed from the hypergeometric distribution using a one-sided Fisher’s exact test and corrected for multiple comparisons using the Benjamini– Hochberg method.

#### Analysis of cell-cell interaction using CellPhoneDB

“CellPhoneDB”^105^ (v5.0; implemented in Python v3.8.19) was used to infer “relevant” differences in cell-cell interactions between Cluster 1 and pro-viral Cluster 2 astrocytes across the scRNA-seq dataset. Specifically, “relevant” interactions were identified from the list of differentially expressed genes comparing Cluster 1 and pro-viral Cluster 2 (p < 0.01; LFC > 1; n= 2,186 enriched in Cluster 1; n= 877 enriched in Cluster 2), a subset of which encode ligands (n=427) or receptors (n=341). “Relevant” interactions were those that were selectively enriched in either astrocyte cluster following the CellPhoneDB gene shuffling approach across all cell-type clusters (neurons, microglia, hNSC and mitotic) from full scRNA-seq dataset (p < 0.05). The “relevant_interactions” CellPhoneDB output file was used for downstream analysis, visualization and plotting in R including, the R package “circlize” (v0.4.16)^123^.

#### Pseudotime trajectory analysis astrocytes using Monocle3

“Monocle3” was used to perform a pseudotime-based trajectory analysis of astrocyte transcriptional diversity following a standard workflow based on six principal components that describe gene expression variation across the human genome. Pseudotime relationships were visualized both through both a UMAP embedding and in a linear fashion after binning scRNA profiles into 40 pseudotime-ordered bins (n=99 or 100 scRNA profiles per bin). Experimental identities associated with the cells in each bin were analyzed as proportions (Cluster 1 and Cluster 2 identities from the Seurat analysis or experimental origin). Viral mRNA load scores were also plotted by bin. To visualize expression signatures of genes induced in pro-viral Cluster 2 (p < 0.01; LFC > 1) across pseudotime bins, bin-specific average expression values were calculated and normalized to maximum expression value to facilitate cross-gene comparisons. Genes were then catergorized by pseudotime expression profile. To do this, each gene’s expression profile was visually inspected. Genes with minimal expression fluctuation before bin 36 yet a clear increase in expression at bin 37 were categorized as “Early”. Genes with minimal fluctuation before bin 36 but were categorized into (mutally exclusive) “Progressive”, “Sustained”, “Transient” or “Late” categories based on expression patterns across bins 38-40.

#### Random forest regression analysis of human gene RNA levels and viral mRNA load

Random Forest regression analysis was performed using the randomForest(v4.7-1.2) R package independently for neurons, Cluster 1 astrocytes and pro-viral Cluster 2 astrocytes. Viral mRNA load scores were used as the dependent variable and normalized gene expression values for those n=11,943 genes that passed expression threshold criteria for differential expression testing using Libra. Regression results were generated from n=200 trees and importance scores were retained using default parameters (*randomForest(ntree=200, keep.forest=FALSE, importance=TRUE, do.trace = 5*).

#### Identification of pro-viral Cluster 2-like Astrocytes in the NPH snRNA-seq dataset

To test for evidence of pro-viral Cluster 2-like astrocytes in the human brain without confounds of post-mortem interval, we took advantage of a published snRNA-seq dataset profiled from flash-frozen cortical tissue acquired during surgical treatment of Normal Pressure Hydrocephalus^79^. We used RNA count data and associated patient-specific metadata from the subset of snRNA-seq profiles belonging to astrocytes (n= 82,634 profiles; https://zenodo.org/records/8319719). The RNA count data were cell-normalized and gene-centered using the standard Seurat workflow (equivalent to the processing performed on the Tha scRNA-seq dataset). To identify potential Cluster 2-like astrocytes from these NPH data, we generated an aggregate expression score based on the n=196 genes observed in the NPH data and enriched in Cluster 2 (p <0.01 and 2 > log2 fold-change). To critically evaluate the biological significance of these aggregate expression scores, we generated 1,000 expression-matched sets of 196 genes. Expression-matching was performed by rank-ordering the NPH genes by expression level; dividing these expression-ranked genes into ten bins from low to high expression; and randomly sampling (without replacement) each bin by the number of instances of each of the 196 Cluster 2 enriched genes. To facilitate comparison of the real and permuted aggregate gene score distributions, each of 82,634 scores were z-scored by gene list and z-score distributions were then analyzed (**Figure 4H**). A z-score threshold of 4.5 standard deviations was used to identify n=38 Cluster 2-like NPH astrocytes contributed by n=21 of 52 total patients (n=1 to 4 Cluster 2-like NPH astrocytes in each of those 21 patients; z-score range: 4.5 – 12.5). The cut-off of 4.5 standard deviations corresponds to a p-value of < 4.7E-7 based on the permuted data. To validate these 38 astrocyte RNA profiles resembled those of Cluster 2, we tested whether gene counts associated with the 196 “discovery” gene set and an independent “validation” gene set of RNAs enriched in Cluster 2 (p <0.01 and 1-2 log2 fold-change; n=521 genes) were enriched in the 38 Cluster 2-like astrocytes versus all other NPH astrocyte RNA profiles (n=82,596). The discovery and validation gene counts were 2.8 and 1.2 enriched respectively in the 38 Cluster 2-like NPH astrocytes and highly significant (chi-square test of expected versus observed UMIs; p < 1E-16).

#### Comparing “reactive” astrocyte gene signatures from other datasets to pro-viral Cluster 2 astrocytes

To assess the similarity of previously reported rodent and human reactive astrocyte gene expression signatures to those observed in pro-viral Cluster 2 astrocytes, we re-analyzed data from *ex vivo* mouse astrocytes stimulated with lipopolysaccharides (Hasel et al; scRNA-seq data)^81^ or cultured rat and human astrocytes stimulated with TNF, IL1α, C1Q, and IFNβ, a cocktail that induces an interferon-responsive reactive state (Prakash et al; RNA and proteomic profiling)^80^. Specifically, for Hasel et al, gene enrichment data from the ten scRNA-seq-defined astrocyte clusters were downloaded (Supplementary Table 5). Analysis focused on those genes enriched in cluster 8 (n=70 genes), an astrocyte sub-population described by Hasel et al as “interferon reactive”^81^. For Prakash et al, RNA and protein enrichments associated with rat and human interferon-responsive state were downloaded (Supplementary Tables 3 and 5). Analysis focused on the n=67 rat genes with concordant, statistically-significant enrichments in both RNA and protein; for human astrocytes, analysis focused on the n=8 genes enriched via proteomic analysis (RNA expression data was not acquired). The genes from these three lists were visualized on volcano plots describing Cluster 1 versus pro-viral Cluster 2 differentially expressed genes. In all three cases, RNAs from the published gene set showed no obvious enrichments for those genes differentially expressed in Cluster 1 or pro-viral Cluster 2 and thus no statistical test was performed.

#### Analysis of single-cell RNA sequencing data

Statistical testing and plot generation was performed in R (v4.3.0; R packages are provided in the Key Resources table). All p-values are reported as adjusted for multiple-hypothesis testing unless otherwise stated. Test-specific information is included the relevant STAR Methods sections.

